# Evolutionary stability of social commitment

**DOI:** 10.1101/2022.07.12.499667

**Authors:** Yuka Shirokawa, Masakazu Shimada, Nao Shimada, Satoshi Sawai

## Abstract

Conflict resolution between individual cells and a group is essential for multicellularity. The social amoeba *Dictyostelium discoideum* switches between solitary growth and social fruitification depending on nutrient availability. Under starvation, cells form fruiting bodies consisting of spores and non-viable altruistic stalk cells. Once cells socially committed, they complete fruitification even with a renewed source of nutrients. This social commitment is puzzling because it deprives individual cells of benefits of quickly resuming solitary growth. One idea posits that traits that facilitate premature de-commitment are somehow hindered from being selected. We studied outcomes of premature de-commitment by forced refeeding. We show that when refed cells resume sociality together with non-refed cells, besides some becoming solitary outside of fruiting bodies, a large fraction was redirected to a sub-region of altruistic stalk regardless of their original fate. The refed cells exhibited reduced cohesivity and were sorted out to the altruistic positions in morphogenesis. Furthermore, a theoretical model considering evolution of cell-cell association revealed a valley in the fitness landscape that prevents invasion of de-committing mutants. Our results provide a general scheme that naturally penalizes withdrawal from a society by evolving a specific division of labor that less cohesive individuals become altruists.

**Significance Statement:** Evolution of unicellular to multicellular organisms must resolve conflicts of reproductive interests between individual cells and the group. In the social amoeba *Dictyostelium*, a transition from a solitary to multicellular group occurs under starvation. Once cells commit themselves to multicellular organization, the process continues even when shifting to an environment that favors solitary growth. Our study revealed that cells forced to partially revert to a de-committed state take an altruistic role through interaction with socially committed cells. The de-committed cells exhibited reduced cohesivity and were sorted out to altruistic positions in morphogenesis. This inevitably penalizes ‘selfish’ cells that revert to solitary growth too quickly. Our results explain group-level behavior that is apparently difficult to understand from an individual-level fitness.

## Main Text Introduction

Cohesion of a group is fundamental to making cooperation fruitful for group members. However, evolution of cooperation is often faced with selection pressures that potentially lead to their collapse. Cheating that exploits benefit of cooperation at the cost of others is a well-studied case of such risks. Nature has found ways to reduce deleterious effects of cheating through interactions among kin [1] and policing that suppress individual proliferation [2]. In microbial cooperation [3], colony expansion that keeps the cooperator cells at close distances [4, 5] and kin recognition using cell-surface proteins [6, 7] play pivotal roles in cheating suppression. Besides cheating, cooperation is also vulnerable to transient environmental changes that raise fitness of a cell as a solitary individual rather than as a social group member. Indeed, evolutionary loss of social behavior has been experimentally demonstrated in myxobacteria under conditions that impede their social interaction while favoring solitary reproduction [8]. Phylogenetic analyses have also shown that the social loss appears to be a common phenomenon across taxa from insects to unicellular organisms [9–11]. Social behavior exemplified by a high degree of division of labor and reproductive skew is thought to have crossed a ‘point of no return’ [12, 13] back to a solitary lifestyle due to its high complexity. The stabilization of cooperation against adverse evolutionary pressures is the key driving force that increases social complexity [14, 15].

How sociality is precluded from reverting to a free-living lifestyle is especially non-trivial in social groups consisting of individual cells that can survive on their own. One explanation is that social traits act disadvantageously for unicellular growth. For example, in the experimental evolution of fission yeast, high rates of apoptosis evolve to promote snowflake-like chains of cells with high nutrient availability. Since apoptosis is disadvantageous for solitary proliferation [14, 16], multicellularity should be maintained [14, 15]. However, such a trade-off may be limited to cases where individual cells can immediately escape the group. The example of the social loss in myxobacteria, which exhibits intermediate social behavior such as reduced sporulation and collective motility [8], hints at the importance of intermediate phenotypes in the evolutionary transition [17]. Given that even simple multicellularity is a multifactorial trait regulated by a large number of genes [18], loss of sociality almost inevitably proceeds through a transition stage where a newly emerging lineage exhibits the intermediate phenotype, meaning that it still interacts with other socially committed cells. In this situation, the social loss may be prevented if the intermediate phenotypes are somehow hindered from being selected, but at least, a new costly countermeasure should not evolve because they are in an environment that favors solitary growth. One of the simplest ways is that the preexisting social defense itself comes with a rather unexpected bonus of acting inhibitory to reversion to a solitary lifestyle. To obtain the unexpected bonus, the defensive function should have incomplete target specificity. For example, social microbes evolve traits that suppress the sporulation of cheaters [19, 20], but how specific it is to a certain genotype is poorly explored. In other examples, bacteriocins spite cells that are sensitive to the signal in an indiscriminatory manner [21]. Given that such elimination of ‘nonself’ is a common and fundamental defense strategy in organisms [22], what may underlie defensive behavior are exclusion of deviators from the original members, which would inevitably suppress the intermediate phenotypes. Taken together, how the intermediate phenotypes between a social and solitary state interact with the original social members is critical in understanding the maintenance of social behavior.

The social amoeba *Dictyostelium discoideum* is a soil-living eukaryote with facultative sociality under non-nutrient conditions [23, 24]. The cells grow and divide in the presence of nutrient sources such as bacteria and fungi, however, when starved, they aggregate to form a multicellular slug that culminates in a fruiting body composed of differentiated spores and stalk cells. *Dictyostelium* has served as a model microorganism to study cellular cooperation [25], since death of stalk cells that support the spores [23, 24] can be viewed as altruistic behavior. Interestingly, after 4 to 6 hours of starvation onward, cells become engaged in the differentiation process [26], and will no longer quickly revert back to the growth phase even with renewed nutrient supply [27]. Instead, cells continue with the formation of fruiting bodies within 24 hours. The inability of differentiating cells to switch back to solitary growth (hereafter referred to as social commitment) is dependent on the multicellular context. When cells that are mechanically dissociated are replenished with nutrients, they revert to solitary growth [26–28]. Thus, unless mechanical dissociation, the only way by which cells return to the solitary state is to mature into spores and then germinate, a long process that takes a few days. Evolutionary benefit of the social commitment is puzzling, especially given that the social amoeba colonizes sites with a frequent supply of rich dung or leaf mold [29], which favor quick reversion to solitary growth. Existence of a phagocytic cell type that removes intruding bacteria within a slug [30] suggests that it is not unusual for socially committed cells to be exposed to nutrients. To understand the multicellular basis that maintains the social commitment in *Dictyostelium*, we studied outcomes of premature de-commitment. In our experiments, cells were forced to take the dedifferentiation path by short-term refeeding, and their fates when mixed with socially committed cells were traced by live-cell imaging. Based on the observation, a theoretical model was constructed to predict the evolutionary stability of the social commitment.

## Results

### Forced refeeding experiments

Mechanically dissociated cells re-plated on agar without nutrients are known to rapidly reaggregate and recapitulate the respective developmental stages within a few hours [31]. We exploited this unique property to investigate the interactions between socially committed and partially de-committed cells. For the de-committed cells, dissociated cells from the slug stage were fed with bacteria *Escherichia coli* B/r for 3 or 5 h (hereafter referred to as RF3 and RF5 cells, respectively), then nutrients were removed again. The refed cells were then allowed to reaggregate with cells that were not refed (NF cells, socially committed cells) (Figure 1*A*). This treatment mimics a situation that a rare cell makes a slug with socially committed cells and dedifferentiates to engulfing bacteria within a slug, where the bacteria remained for 3 to 5 h before the terminal differentiation (18 to 23 h after starvation). A strain harboring the prestalk (ecmAO-RFP) and prespore (pspA-GFP) marker genes were used for identification of their original cell fate owing to the time for degradation of fluorescent proteins (> 4–5 h) [32]. The mixture of cells jointly formed a tight mound after 1.5–2 h (Figure S1, Movies S1 and S2). In the mixture of NF cells with and without the cell-type reporter (NF+NF), prestalk cells were found in the peripheral region of the aggregate, while prespore cells were more uniformly distributed (Figure 1*B*: NF+NF) in line with the previously described sorting pattern [33]. In contrast, in the mixture of RF and NF cells, RF cells (fluorescent cells) were positioned at the periphery regardless of the original cell type (Figure 1*B*: RF3/5+NF), while NF cells (non-fluorescent cells) were more abundant in the aggregation center. Thus, the refed treatment significantly affected cell positioning (Figure 1*C*, Table S1: overall model: GLM and analysis of deviance, cell fate: *P* < 0.0001, treatments: *P* < 0.0001, prespore cells in NF+NF versus RF5+NF: *t*-test, adjusted *P* < 0.0001). In the swap control, we observed a consistent pattern (Figure S2*A*, RF(Fl-)+NF(Fl+)), indicating that expression of the marker genes itself did not affect the sorting pattern. Furthermore, our supporting data indicates that the segregation was relative behavior when RF cells interact with the NF cells because RF cells were still fully capable of developing on their own (Figure S2*B*), and the unique sorting pattern was not merely due to mechanical interruption but also required nutrient replenishment (Figure S2*C*).

**Figure 1.**
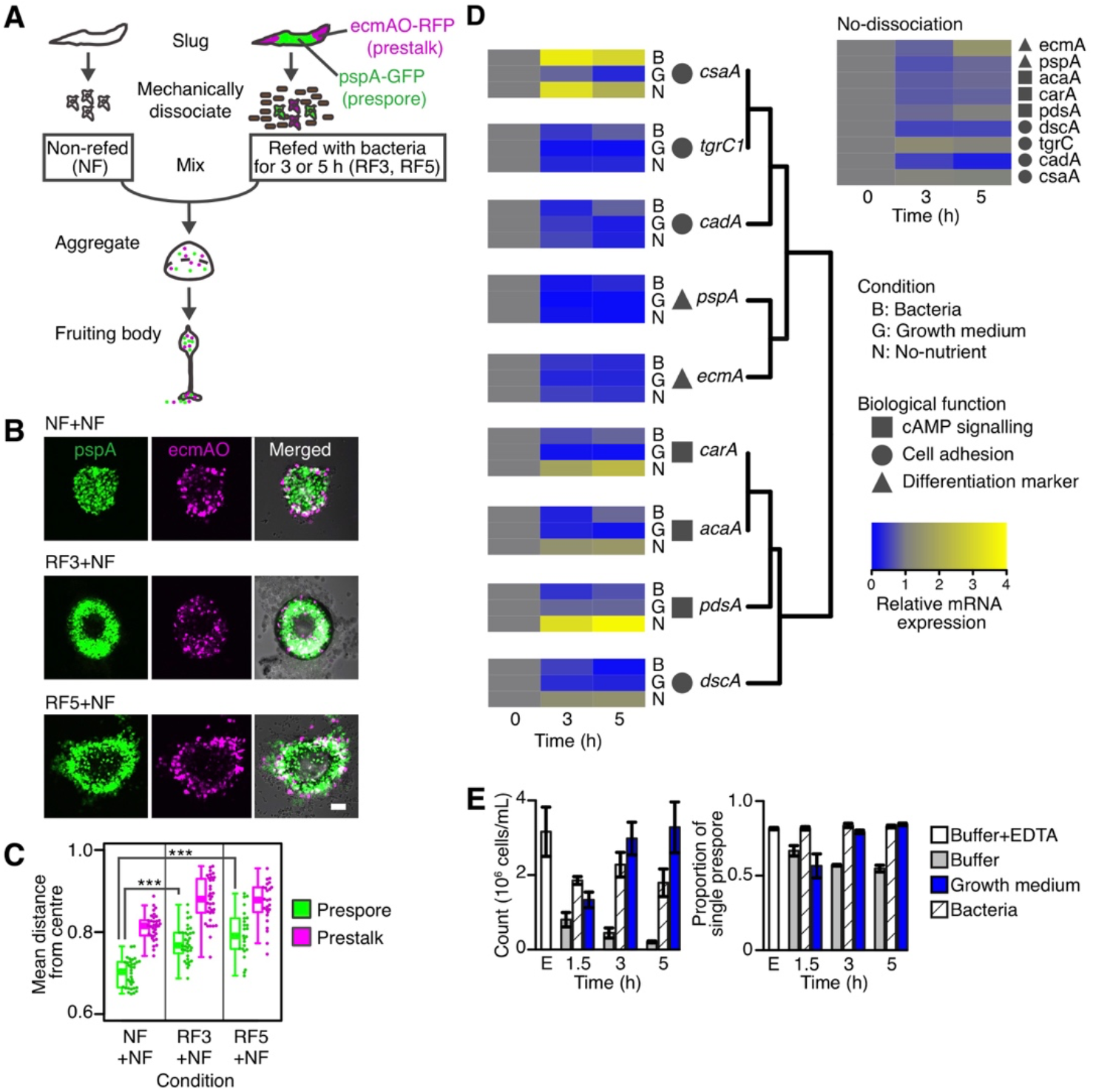
Prestalk-like positioning and cell-state of refed cells. (*A*) A schematic of refeeding and reaggregation of a cell mixture. Dissociated cells were shaken with *E. coli* for 3 or 5 h (Refed: RF3, RF5), washed and mixed with non-refed cells (Non-refed: NF). RF cells harbored prespore (pspA-GFP, green) and prestalk (ecmAO-RFP, magenta) markers. (*B*) Representative images of mixed aggregates 1.5–2 h after plating (pspA: GFP-channel, ecmAO: RFP-channel, Merged: bright-field and fluorescence images). ‘NF+NF’: a mixture of NF cells with and without the cell-type reporter as a control condition. ‘RF3+NF’ and ‘RF5+NF’: Mixture of NF cells and RF3 or RF5 cells, respectively at RF:NF = 15:85 ratio. A scale bar = 50 μm. (*C*) Normalized mean distance between the aggregate center and cells of prestalk- or prespore-origin. The sample size N (biological replicate, aggregates): NF+NF = (4, 34), RF3+NF = (4, 35), RF5+NF = (3, 25). Circles: Values for individual aggregates, lines inside of box plots: The median values. *** P < 0.0001 by *t* test. (*D-E*) Cell-state change incurred by refeeding. (*D*) qRT-PCR analysis. Gene expression levels of dissociated cells in bacterial suspension ‘B’, growth medium ‘G’, non-nutrient buffer ‘N’ (left panel). Unperturbed slugs 18 h into starvation onward (‘No-dissociation’: Upper right panel). Gene expression levels were normalized by the 0 h point. Gene targets were clustered hierarchically. The sample size N = 3 are biological replicates. (*E*) Differences in cell cohesiveness between RF and NF cells. The number of single cells within a sample (left panel). The proportion of single prespore cells in the total single cells within a sample (right panel). ‘E’: Cells in the phosphate buffer containing EDTA (Buffer+EDTA) as a control (most cells were in a single state). The number (1.5, 3, and 5) indicates hours shaken in suspensions. The sample size N =3 are biological replicates, with 380–5966 cells per condition. Error bars are standard error.

Quantitative real-time polymerase chain reaction analysis (real-time PCR, Table S2, see *SI* Methods) showed that RF cells exhibited reduced expression of cell-type differentiation marker genes (Figure 1*D*, *pspA, ecmA*), indicating partial dedifferentiation (Figure *1D*, Table S1: bootstrap test, adjusted *P* > 0.025). Whereas genes encoding the main adhesion molecules [34] (*cadA, csaA*, and *tgrC1*) did not show nutrient-specific changes, *carA, acaA, pdsA*, and *dscA* genes (Figure 1*D*) showed contrasting activity of being reduced by refeeding and induced in non-nutrient buffer (Figure 1*D*, Table S1: GLM and analysis of deviance, ‘No-nutrient’ versus ‘Bacteria’: adjusted *P* < 0.05, ‘No-nutrient’ versus ‘Growth medium’: adjusted *P* < 0.01). *CarA, acaA*, and *pdsA* are essential for the cAMP relay during cell aggregation, and *dscA* (discoidin I) is implicated in the cell-substratum adhesion during aggregation [35, 36].

Given that RF cells show reduced expression of the re-aggregation genes, they are likely to fall behind in joining cell clusters by delayed chemotaxis. However, RF cells appear well mixed and then sorted out afterward (Figure S1, Movie S2). Although a recent work suggested a sorting mechanism of a tip-forming prestalk subtype [37], RF cells are sorted to the periphery and then to the base of the mound instead of the apical tip (Figure S3*A*). These observations point to an alternative mechanism based on differential adhesion [38], where cell-type that position themselves in the peripheral region of a cell mass should be less cohesive than those positioned in the inner region. We tested the ability of RF cells to associate with aggregates using a cohesion assay [39]. Briefly, a suspension of mechanically dissociated cells was shaken to facilitate cell agglutination for 1.5, 3, and 5 h, and the number of single unattached cells was quantified (Figure *1E* left, see SI Methods). We found that cells in the bacterial suspension or growth medium were less cohesive (Figure 1*E* left) than those in the phosphate buffer (Figure 1*E* left, Table S1: GLM and analysis of deviance, Buffer 5 h versus Bacteria 5 h: adjusted *P* < 0.0001, Buffer 5 h versus Growth medium 5h: adjusted *P* < 0.0001) as expected [40]. In addition, we found that the ratio of prespore cells in the unattached cells within the phosphate buffer (55%) was lower than that in the non-adhesive control (82%) (Figure 1*E* right), indicating that non-refed prespore cells are cohesive than prestalk cells in line with earlier literature [41]. On the other hand, refed prespore cells were less cohesive comparable to that of the non-adhesive control (Figure 1*E* right, Table S1: GLM and analysis of deviance, Buffer+EDTA versus Bacteria 5 h: adjusted *P* = 0.572). The lower cohesiveness of prestalk cells relative to prespore cells (Figure 1*E* right) and comparably reduced cell-cell cohesivity in refed cells (Figure 1*E* left) are consistent with the peripheral positioning (Figure 1*B,C*).

Positional bias was also evident during the culmination (4–6 h after plating) in the stalk region. In addition to the main stalk, stalk cells of *D. discoideum* constitute three distinct cell types: The upper and lower cups and basal disc (Figure 2*A*) [24]. The cell mass elongated into a finger-like form with a prespore cell mass between the upper and lower cups (Figure 2*B,C*, Figure S3*B*, Movie S3), and the normal pattern was observed in NF cells (Figure 2*B*: NF+NF). In contrast, in RF+NF populations, both prespore and prestalk cells of RF cells were abundant in the lower cup region (Figure 2*B*, Figure S3*B*, Movie S4: RF3+NF, RF5+NF). Accordingly, the positional frequency of prespore and prestalk cells was anti-correlated in NF+NF, and positively correlated in RF+NF (Figure 2*B*, Table S1: GLM with gamma error, NF+NF: slope = −2.03 ± 0.26, RF5+NF: slope = 3.70 ± 0.31). Later, RF cells formed the lower cup and basal disc but were excluded from the upper cup and main stalk (Figure 2*C*, RF3+NF). In contrast to our standard condition (RF cells were the minority; Figure 2*C*, RF3+NF), when RF cells were the majority, refed prespore and prestalk cells were positioned according to their original cell fate (Figure 2*C*, RF3(Fl+):NF(Fl-) = 70:30).

**Figure 2.**
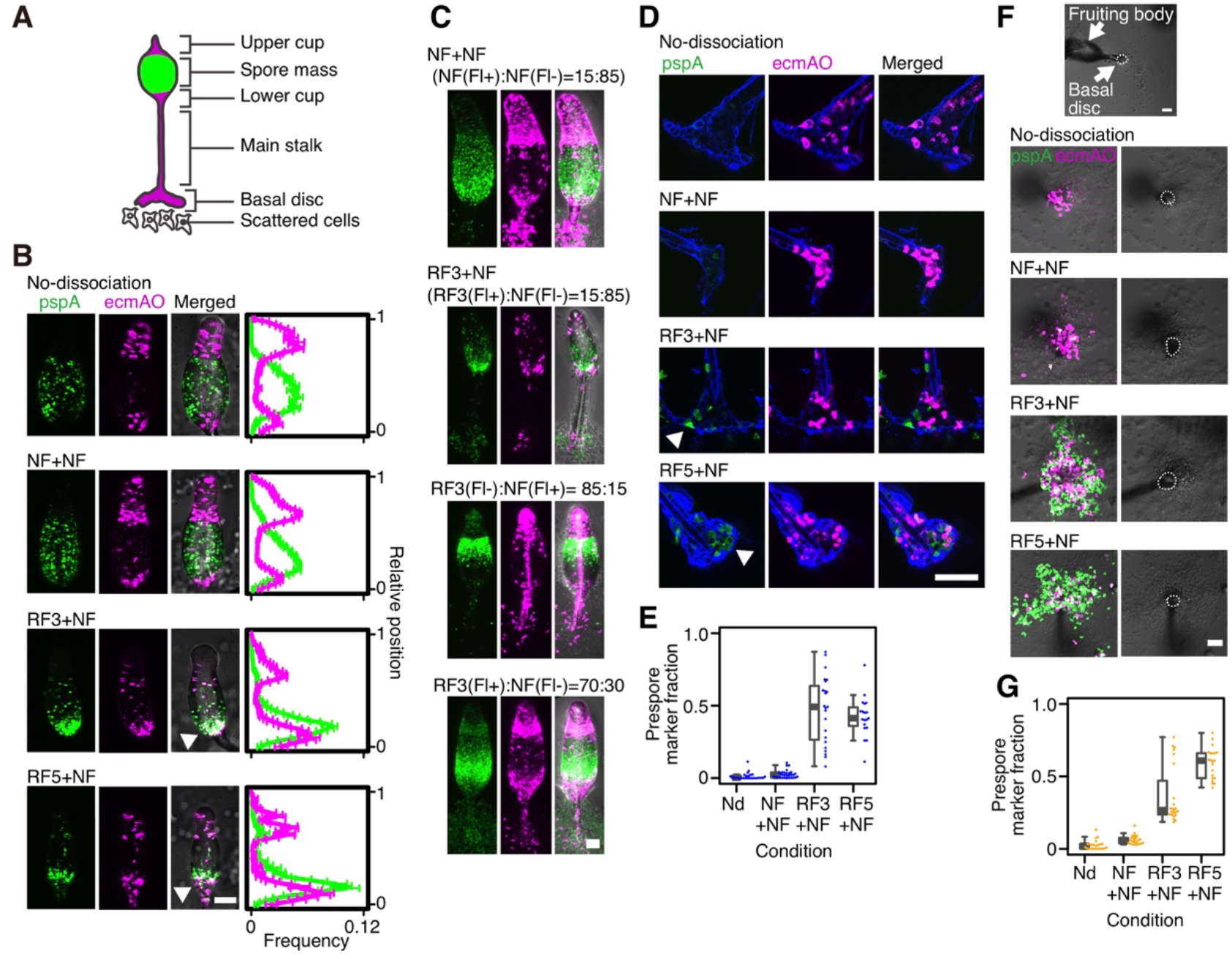
Refeeding redirects cells to the role of prestalk subtype. (*A*) A schematic illustration of the fruiting body. (*B*) Snapshots of the upper region of early fruiting bodies (left). Frequency of prestalk (magenta line) and prespore (green) marker-positive pixels along the anterior-posterior axis (right). No-dissociation (Nd): A fruiting body formed by unperturbed cells. In RF3+NF and RF5+NF, RF cells were found exclusively at the lower cup (arrowheads). The sample size N (biological replicate, fruiting body number): N: Nd = (2, 14), NF+NF = (3, 24), RF3+NF = (3, 20), RF5+NF = (3, 13). For abbreviations, see also Figure 1. (*C*) Early fruiting bodies consisted of different ratios of RF3 and NF cells. Fl+, Fl-: with/without cell-type markers, respectively. (*D*) Fluorescent images of the basal disc. Images of stalk cellulose staining (blue, calcofluor white) were merged with fluorescence images (pspA, ecmAO). Note RF cells of prespore origin are found in the basal disc (RF3+NF and RF5+NF) (arrowheads). (*E*) The ratio of prespore marker-positive to prestalk region in the basal disc. N: Nd = (3, 26), NF+NF = (3, 30), RF3+NF = (3, 23), RF5+NF = (3, 19). (*F*) Representative images of the basal region of fruiting bodies. Note RF cells (fluorescent cells, left panel) scattered around the basal discs (white circles, right panel) in RF3+NF and RF5+NF. (*G*) The ratio of prespore marker-positive region to prestalk region in the scattered cells. N: Nd = (3, 23), NF+NF = (3, 23), RF3+NF = (3, 23), RF5+NF = (3, 21). All scale bars = 50 μm. Error bars: Standard error.

In the terminal differentiation (8 to 10 h after plating), the ectopic positioning of RF cells was most notable in the basal disc region, which consists mainly of secreted cellulose and vacuolating non-viable cells [42, 43]. In the controls (‘No-dissociation’, NF+NF), the basal disc was occupied by cells of prestalk-origin (Figure 2*D,E*) as expected [44]. On the other hand, in RF+NF (Figure 2*D,E*), RF cells of both prestalk and prespore origins were found at the basal disc (Figure 2*D,E*, Table S1, NF+NF versus RF5+NF: *t*-test, adjusted *P* < 0.0001). In addition, cells scattered around the base [45] were amoeboid and showed random movement (Movie S5), indicating escape from death as stalks. In the controls, these scattered cells were of prestalk origin (Figure 2*F,G*: ‘No-dissociation’, NF+NF), whereas in RF+NF (Figure 2*F,G*: RF3+NF, RF5+NF), RF cells of both prespore and prestalk origins were found (Figure 2*F,G*, Table S1: NF+NF versus RF5+NF: *t*-test, adjusted *P* < 0.0001). At the terminal differentiation, a higher proportion of RF cells were allocated to the solitary amoeboid cells rather than spores, compared to NF cells in the mixed cell population of RF+NF (Figure S4). These results (Figure 2*F,G*, Figure S4, Movie S5) indicate that the final survivors at the terminal phase were solitary cells (amoeba) and social cells (spores).

### The growth cost of the social commitment

Our refeeding experiments indicate that *Dictyostelium* cells that dedifferentiate precociously are penalized by being assigned the role of the non-viable stalk (Figure 1,2). Here, we went on to investigate a basic question of how the social commitment imposes individual cells growth costs. Cells were released from the commitment by mechanical dissociation and two types of growth costs were estimated: Growth delay incurred by fruiting body formation (d) and lag time for spore germination (λ) (Figure 3*A*). We found no significant growth differences due to the growth delay *d* (‘Refed & No-dissociation’ versus ‘Refed & Dissociation’: GLM and analysis of deviance, *P* = 0.069); however, as the lag time for spore germination (*λ*), the socially committed cells consumed approximately 9 h longer time to start cell growth than the de-committed cells (Figure 3*B*, Table S1: Estimated lag time between cell inoculation and initiation of cell growth, ‘Refed & Dissociation’: 15.24 ± 4.27 h, ‘Refed & No-dissociation’: 23.97 ± 4.35 h). The significant growth delay highlights the cost for individual cells of remaining in the social group under nutrient-rich environments (Figure 3*B*, Table S1: ‘Refed & No-dissociation’ versus ‘Refed & Dissociation’: GLM and analysis of deviance, *P* < 0.003).

**Figure 3.**
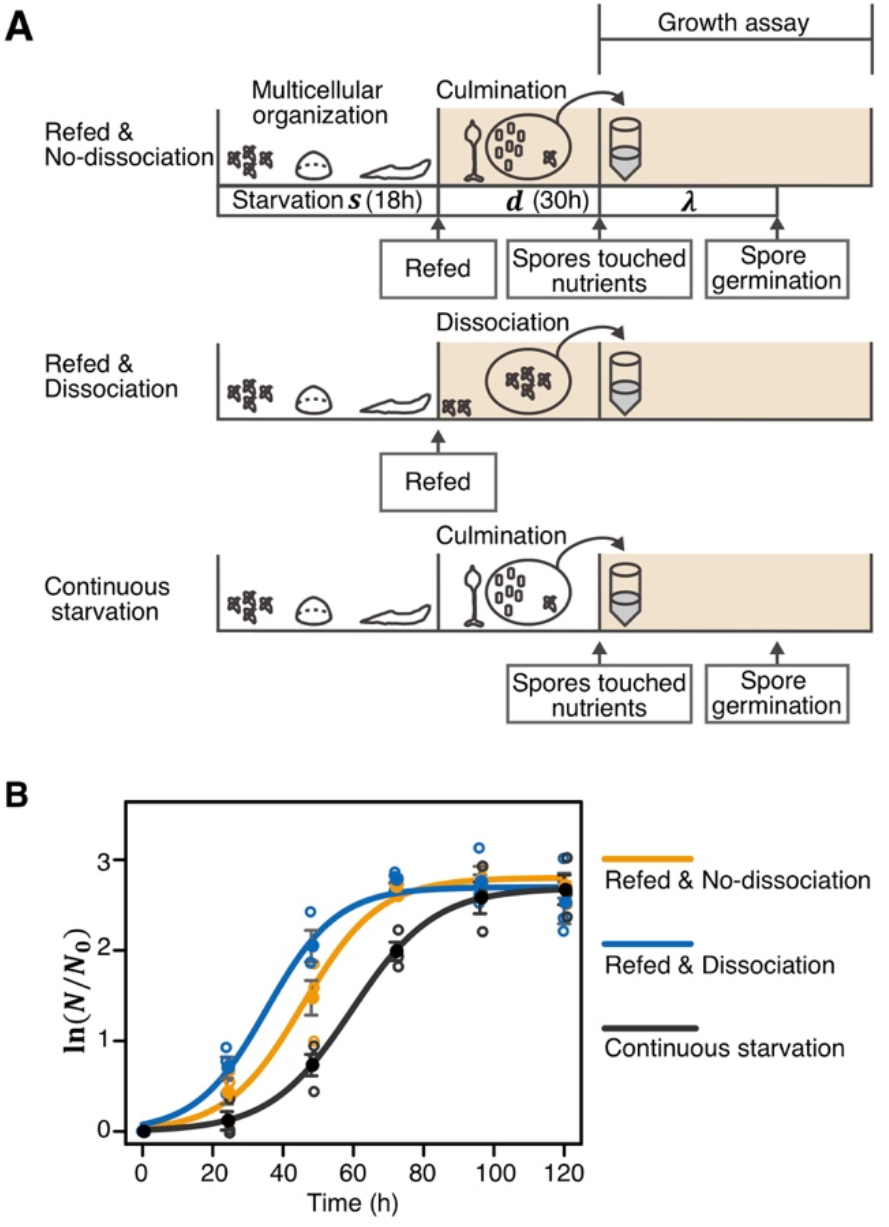
Social commitment imposes growth costs under nutrient recovery. (*A*) The schematic of the growth assay. Cells in the slug stage (s =18 h) are exposed to nutrient recovery. Cells were released from the commitment by mechanical dissociation. Under the social commitment (‘Refed & No-dissociation’ and ‘Continuous starvation’), cells ignore nutrients and develop into a fruiting body. Thus, except for only rare non-aggregated cells, the socially committed cells would bear two types of growth costs: Growth delay incurred by fruiting body formation (d) and lag time for spore germination (*λ*). Without social commitment (‘Refed & Dissociation’), cells quickly restart growing immediately after refeeding. After 30h incubation (d) for the fruiting body maturation, amoeboid cells and spores were harvested and transferred into the growth medium to estimate growth costs. (*B*) Growth assay. ‘Refed & No-dissociation’ (orange): A growth curve of cells from the non-dissociated refed slugs, ‘Refed & Dissociation’ (blue): Cells from refed slugs that were dissociated, ‘Continuous starvation’ (black): cells from uninterrupted slugs under continued starvation. In(N/N_0_): Logarithm of relative population size that was normalized by the initial cell number. The error bars: Relative error. Circles: Individual values from three biological replicates. Regrowth of refed cells from non-dissociated slugs was significantly delayed compared to that of dissociated slugs (GLM, *P* < 0.003).

### Model analysis for evolutionary maintenance of the social commitment

If the de-committed strain is rewarded by such a growth advantage (Figure 3), how stable is social commitment against adverse evolutionary pressures? We analyze whether a rare mutant cell that reverts to a solitary state could invade a wild-type population with the framework of adaptive dynamics [46, 47]. We considered a sufficiently large population of haploid and asexually reproducing cells that are exposed to nutrient recovery at each slug stage. The evolutionary trait is a degree of association with other cells, *x* (0 ≤ *x* ≤ 1) (Table S3). Cells with any of the traits normally commit to the differentiation, and then express the trait *x* at the slug stage. Both cell fate and fitness are decided according to *x*. A wild-type cell has the trait *x* = *x_wt_*, where its default value is *x_wt_ = 1* indicating full social commitment. According to the value of *x*, a single cell simultaneously has probabilities to be a spore, stalk, and solitary cell. The probabilities are equivalent to cell fate ratios in a clonal population with the trait *x*. The cell fate ratios of the full social commitment (*x =* 1) are designated as spore: *SP_initial_ =* 0.6, stalk: *ST_initial_* = 0.2, and solitary: *Sol_initial_* = 0.2, respectively (for suitability of the parameters see, Figure S5).

We consider that invasion of a rare mutant cell with the trait *x*’ = *x_wt_* + *ε*, (*ε* is a sufficiently small negative value). The rare mutant cell makes a chimera slug with the wild-type cells, and expresses the trait *x’* within the slug and reverts to a solitary state. We assume that *x* controls most of cohesion proteins as a ‘master regulator’, i.e., *x* = 1 indicates fully cohesive, and *x* = 0 is complete degradation of cohesion proteins (Figure 4*A*). According to previous study and our experimental results (Figure 1*E*) [41], cohesion of each cell fate is assumed to be distinctively different, and the highest is prespore, followed by in the order of prestalk, solitary cell (hereafter the allocation is referred to as ‘canonical allocation’). When a cell has a reduced trait value (*x* < 1), the trait expression may induce cohesion degradation as the conceptual illustration of distribution shifts in Figure 4*A*. For example, the prespore cell after the trait expression will exhibit reduced cohesion similar to an original prestalk or solitary cell. From our refed experiments (Figure 1,2), a cell is assumed to play a cell fate role as determined by its own reduced cohesivity, rather than its original fate. Thus, given the order of cohesion (Figure 1*E*) [41], a cell changes a role via evolution *x* in the order of prespore, prestalk, and solitary cell (Figure 4*B*). With these evolutionary steps being described by the following differential equations:

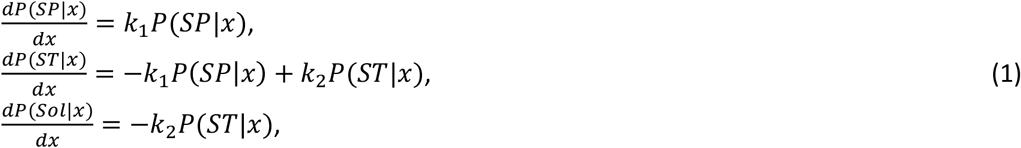

where *P*(*Z|x*) is the probability of a cell with the trait *x* taking fate *Z* (SP: Spore, ST: Stalk, Sol: Solitary cell). Note that *P*(*SP*|*x*) + *P*(*ST*|*x*) + *P*(*Sol*|*x*) = 1. *k*_1_, and *k*_2_ are the rates of cell fate transition (*k*_1_ > 0, *k*_2_ > 0). If *k*_1_ ≫ *k*_2_, decrement of the spore probability is quick but that of stalk probability is slow in the evolution (Eq. **1**, Figure 4*B*). On the other hand, if *k*_1_ ≪ *k*_2_, vice versa. Our refed experiments (Figure 2) support the assumption *k*_1_ ≫ *k*_2_, because we observed a much smaller number of RF cells in the prespore region of early fruiting bodies (quick *P*(*SP*|*x*) decrement) but abundant RF cells in the stalk region (lower cup and basal disc) (slow *P*(*ST*|*x*) decrement). We select default values as *k*_1_ = 14.6 and *k*_2_ = 6.3 (see, SI text).

**Figure 4.**
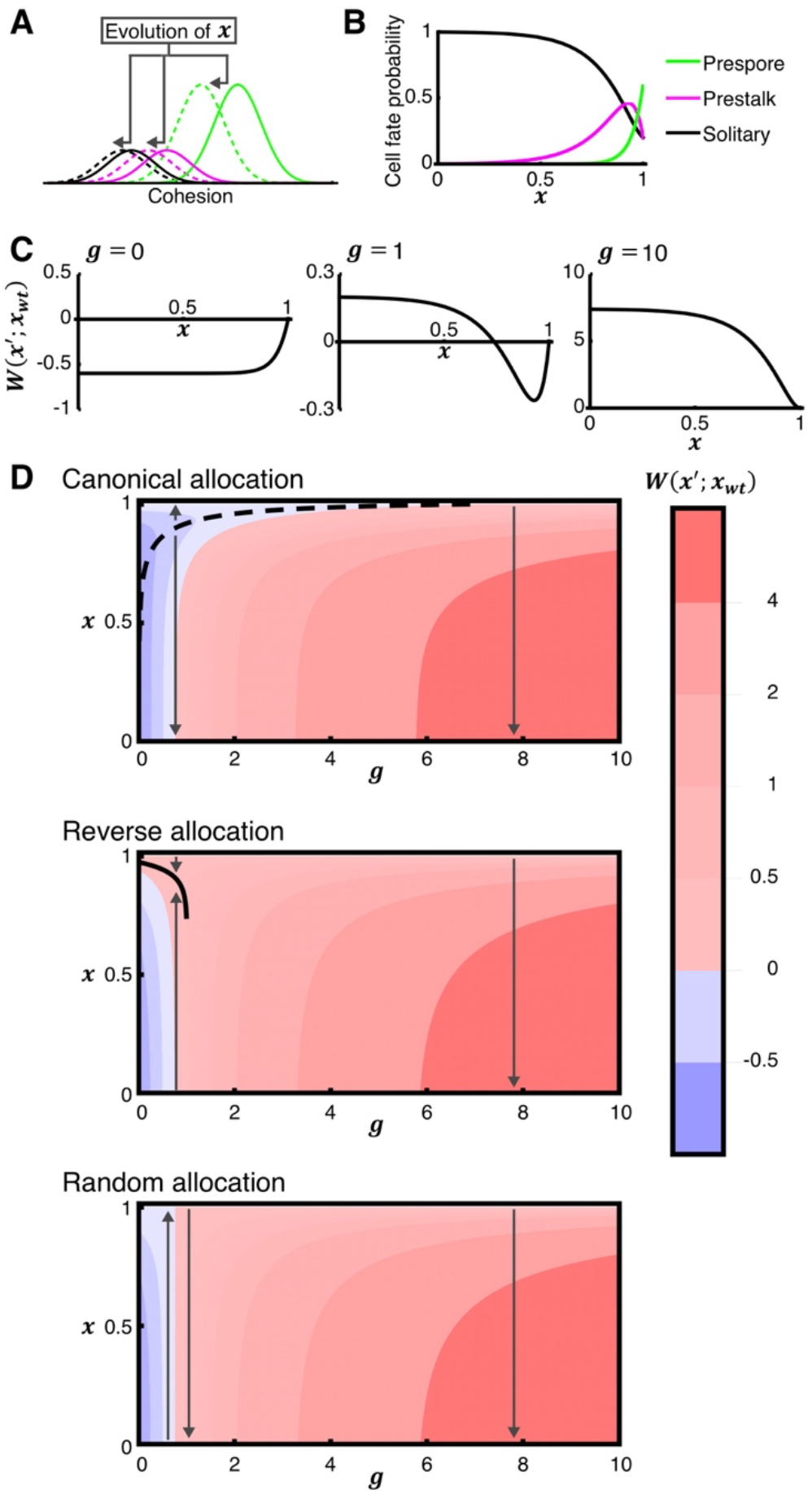
Social commitment in *Dictyostelium* is an evolutionarily stable strategy. (*A*) An intuitive explanation of cell fate transitions via the evolution of trait *x*. The trait *x* (0 ≤ *x* ≤ 1) is assumed to decide the cell-cell association level, which generally controls most of cohesion proteins as a ‘master regulator’ (*x* = 1: full social commission, *x* = 0: a solitary state, complete degradation of cohesion proteins). The solid lines indicate cohesion distributions of prespore (green), prestalk (magenta), and solitary cells (black) in the cell population. The evolution of reduced *x* can be illustrated as distribution shifts (dotted line). After evolution of *x*, prespore cells (dotted green line) will exhibit reduced cohesion similar to original prestalk and solitary cells (solid magenta and black lines). Cells are assumed to play a cell-fate role as determined by the reduced cohesion rather than the original cell fate, according to our experiments (Figure 1,2). (*B*) Evolution of *x* and its link to the cell fate role probability assuming division of labor that assigns each cell fate with a degree of cohesion (canonical allocation: prespore > prestalk > solitary cell) (Eq. **1**). (*C*) The invasion fitness of the solitary reversion mutant within the wild-type population *W*(*x’*; *x_wt_*), where the wild-type trait is *x_wt_* = 1 under canonical allocation. *g* is the additional weight for the solitary fitness as defined by nutrient availability. Note that the fitness function has a valley in the intermediate nutrient condition (*g* = 1) suggesting that the mutant cannot evolve when the evolution starts from the vicinity of *x* = 1. (*D*) Evolutionary directions and equilibrium under various environmental conditions. Black lines indicate analytical predictions for the singular point *x\** (dotted line: Evolutionary repellor, solid line: Evolutionary attractor). Allows indicate the direction of selection. The red and blue colors indicate invasion fitness of the mutant interacts with the wild-type cell with *x_wt_ =* 1.

Invasion fitness of the mutant cell is

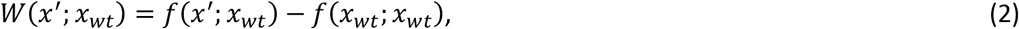

where, *f*(*x’*; *x_wt_*) is the survival probability of the mutant, and *f*(*x_wt_*; *x*_wt_) is that of the wild-type cell (Table S3). Both *f*(*x’*; *x_wt_*) and *f*(*x_wt_*; *x_wt_*) are defined as the sum of the probabilities of becoming a spore (social fitness) and a solitary amoeboid cell (solitary fitness) as follows,

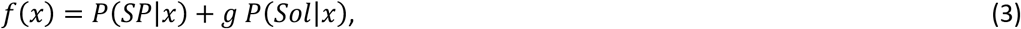

where, *g* represents the additional weight of solitary fitness as defined by amounts of nutrients. Thus, for simplicity, amounts of nutrients directly contribute the solitary fitness. The equation Eq. **3** simply describes the allocation between solitary and social life, and we also consider another formulation that assumes explicit altruism (Figure S6) shown fundamentally the same result as that from Eq. **3**.

The invasion fitness when the mutant interacts with the wild-type cell with *x_wt_* = 1 (Figure 4*C*) showed a fitness valley under the intermediate nutrient-rich condition (e.g., *g* = 1), owing to the transient increase in the stalk fate probability during the evolution. In the starved condition (*g* = 0), the mutant is always disadvantageous, while in the nutrient-rich condition (*g* = 10) the mutant rapidly evolves (Figure 4*C*). Ecology-evolution-environment diagram [47] in Figure 4*D* shows that the evolutionary equilibrium is *x* = 0 or *x* = 1 depending on the starting point of the evolution. The point where the selection gradient vanishes, i.e., ‘evolutionarily singular’ [46, 47] *x\** is in the ‘bottom’ of the valley (evolutionary repellor) (Figure 4*D*, ‘Canonical allocation’), and neither an evolutionarily stable strategy (ESS) [48] nor convergence stable [46, 49] under our assumed condition (*k*_1_ ≫ *k*_2_) (for detail, see SI text and Figure S7). Thus, the model predicts that once the wild-type cells acquire complete social commitment (*x* = 1), mutant invasion is blocked under a wide range of intermediate nutrient-rich conditions.

The fitness valley (Figure *4C,D*) was caused by a direct consequence of division of labor that assigns each cell fate with a degree of cell-cell cohesion in the order of prespore > prestalk > solitary cell (canonical allocation). If the social amoeba were to have an alternative type of allocation rule: With cohesion order of prestalk > prespore > solitary cell (reverse allocation), and with prespore = prestalk > solitary cell (random allocation), these systems did not prevent invasion of the solitary reversion mutant (Figure 4*D*). In the reverse allocation system, an evolutionary singular point *x*\* is the partially solitary state (Figure 4*D*, ‘Reverse allocation’) and is ESS and convergence stable under our assumed condition (*k*_1_ ≫ *k*_2_) (see SI text), indicating that cells make more spores by the solitary reversion in exchange for small amounts of robustness of a fruiting body. In the random allocation system, there is no singular point (Figure 4*D*, ‘Random allocation’). Taken together, only the canonical allocation system can protect the social commitment under nutrient-rich conditions.

Finally, to understand the range of the social defense, invasion of a cheating mutant is investigated in addition to the solitary reversion mutant. We consider a rare cheating mutant cell that has a higher probability to be a spore than the wild-type cell within a chimera. An evolutionary trait *i* is additional spore investment at the slug stage. The wild-type trait is *i* = *i_wt_* where *i_wt_* = 0 is the default value, and the cheating mutant trait is *i* = *i*’ = *i_wt_* + *θ*, (*θ* is a sufficiently small positive value). We especially focused on the detailed developmental process of the social amoeba: a slug migrates toward the light, probably to reach the soil surface, where spore dispersal is more likely [24]. The migrating slug induces a series of cell fate transitions closely resembling those associated with the canonical allocation: There is a sub-population of prestalk cells left behind from a slug [50], which are compensated by trans-differentiation of prespore to prestalk cells [51]. Cell fate transition over the migration time *t* is described as differential equations (Eq. **S17**-**S19** in SI text). The invasion fitness of the cheating mutant *W*(*i*’; *i_wt_*) is the sum of spore and solitary probabilities (Eq. **S21**, Eq. **S22** in SI text). The selection gradient (Eq. **S23** in SI text) was constant with respect to _i_, and we found the condition that the cheating mutant evolves (red lines in Figure 5, Eq. **S24-S26** in SI text). The model shows that evolution of the cheating mutant is suppressed under nutrient-rich conditions in the canonical allocation, while in the reverse allocation the evolution is suppressed under starved conditions but enforced under nutrient-rich conditions. In the random allocation, cheating always evolves (Figure 5). Thus, if the social amoeba experiences only starved conditions, the reverse allocation is suitable; however, under nutrient-rich conditions, the canonical allocation will demonstrate the comprehensive defense capacity.

**Figure 5.**
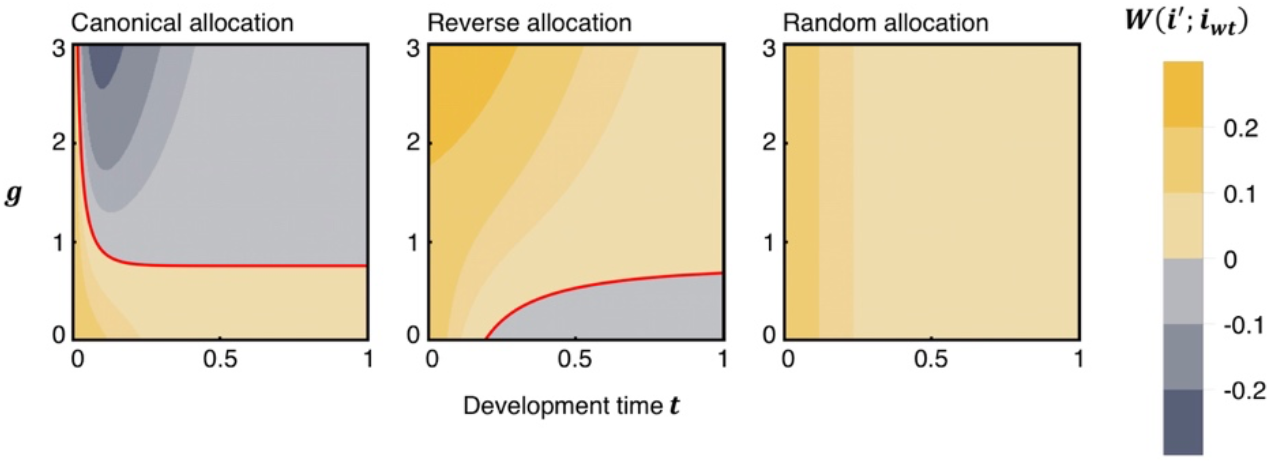
Division of labor in *Dictyostelium* effectively reduces cheating advantage. Evolutionary trait *i* describes additional spore investment, *i_wt_*: The wild-type cell trait, *i*’: The cheating mutant trait (*i_wt_* < *i’*). *W*(*i’*; *i_wt_*): Invasion fitness of the cheating mutant within the wild-type population. *t*: development time with slug migration. *t* = 0: The slug stage, *t* = 1: the maximum migration time. Red lines indicate the analytical predictions of whether the cheater invades the wild-type population. The cheater can invade under the conditions below the red line in the canonical allocation system, and above the line in the reverse allocation system. In random allocation, cheating always evolved. Initial cell fate probability of the mutant being prespore: 0.8, prestalk: 0, solitary cell: 0.2, that of the wild-type strain being prespore: 0.6, prestalk: 0.2, solitary cell: 0.2 are shown.

## Discussion

Maintenance of cooperation is inextricably linked to the choice of social members to be dispersed from a society [52]. We show the choice of the leavers can canalize ‘point of no return’ in the social evolution [12, 13]. Our results suggest that *Dictyostelium* employs the division of labor in which cells with a less cohesive propensity become altruists. The dedifferentiating *Dictyostelium* cells take a prestalk-like cell state through interaction with differentiating cells (Figure 1,2); thus, the system penalizes precocious solitary reversion (Figure 4). Unlike policing, which entails inescapable costs in general [2], the penalization occurs naturally through fruiting body formation and the social amoeba seems not to pay explicit costs.

What evolutionary trajectory did organisms follow to cross ‘point of no return’ is one of fundamental questions. In addition to the solitary reversion mutant, our model (Figure 5) showed that the division of labor of *Dictyostelium* can effectively reduce the advantage of cheating mutants, which is consistent with earlier experimental observations [53]. From these results, it is tempting to speculate that the social amoeba could have first acquired the canonical allocation to suppress cheating, and this in turn penalized solitary reversion as a byproduct. Given that the dedifferentiation process is regulated by several thousand genes [28, 54], the presence of the fitness valley (Figure *4C,D*) acts as a strong resistance to the social commitment breaking down. Furthermore, for microbes with large clonal populations, the chance of crossing the fitness valley through genetic drift is slim [55] without additional mechanisms [56]. On the other hand, the fitness valley may work only at the early stage of invasion due to its frequency dependency; The de-committed cells were penalized when they are in the minority (~15% of the population) but not in the majority (70%~) (Figure 2*C*). Even if the social amoeba crossed the valley, the evolutionary progression may be too slow to manifest itself, since the dedifferentiation process is known to be robust to genetic perturbation [54]. Future experimental evolution studies should clarify the evolutionary process given their genetic background and developmental processes.

How the social commitment of *Dictyostelium* is maintained has been a long-standing question ever since the observation was first reported [27]. Our experimental results suggest an interesting cellular dynamics-based mechanism as to how this is implemented. The initial prestalk-like positioning of refed cells to the mound periphery (Figure 1*B*) can be explained based on the mechanism of cell segregation [38] by differential adhesion of non-refed cells and refed cells (Figure 1*E*). Later, the refed cells were positioned at the base of the mound (Figure S3*A*), which then formed the lower cup (Figure 2*B*) and the basal disc (Figure 2*D*). The basal disc and lower cup are well-known terminal states of prestalk cells [44, 57], however, how cells are positioned at these locations is not well understood. The localization of refed cells is distinct from that of the primary stalk cells but similar to that of the ‘anterior-like cells’ [57–60] – a small percentage of poorly characterized prestalk subtype that constitutes the basal disc, the lower cup, and the upper cup. The lack of refed cells from the upper cup makes it distinct from the anterior-like cells. This, as well as its absence from the main stalk, may be related to reduced sensitivity to cAMP due to the decreased *carA* gene expression in the refed cells (Figure 1*D*), as low sensitivity to cAMP facilitates re-differentiation of prespore cells to lower cup cells and inhibits prestalk cells positioning to the upper cup [61].

Cooperation is a fundamental force that leads to the emergence of collective-level selection units [62, 63]. Paradoxically, the fact that individuals are allowed to choose a solitary strategy among other options, such as to cooperate or to cheat, can in turn maintain cooperation by reducing cheating interest [64]. In cellular cooperation, the presence of cells ungrouped can compensate for the cost of aggregation [65], and facilitate coexistence of diverse fruitification traits as a bet-hedging strategy to variable environmental conditions [66, 67]. However, such a solitary cell type may not be sufficient to maintain cooperation in environments that act favorably for solitary growth (Figure 3*B*), for example, relatively long nutrient-rich conditions interrupted by brief intervals of starvation. Under such conditions, precocious reversion to a solitary state would be beneficial for individual cells. This notion is supported not only by our growth assay (Figure 3*B*) but also by the fact that there exist cells that are left behind from a slug that can grow quickly after nutrient recovery [68]. The social system appears to include a certain proportion of a solitary cell type in the first place as a bet-hedging strategy, but for those that are socially committed, there is little payoff from quitting cooperation because of their specific division of labor (Figure 4). Our study provides a simple explanation for the emergence of collective behavior that is apparently counterintuitive in terms of individual-level fitness.

## Materials and Methods

### Refeeding experiments

Strains expressing cell fate markers were constructed from AX4 cells. To obtain slug-stage cells, vegetative cells were washed with the phosphate buffer (PB) and applied to a 1% PB agar plate and incubated for 18 h. PB containing 20 mM EDTA was applied to the slugs, and the cells were mechanically dissociated by repeated pipetting and passed through a 23 G needle and 40 μm cell strainer. For refeeding, dissociated cells were co-suspended at 2 to 3 × 10^6^ cells/mL with the concentrated *E. coli* B/r. NF cells were cells suspended in PB immediately after dissociation. The mixed cell suspension contains RF and NF cells at 1 to 2 × 10^6^ cells/mL was plated on a glass-bottomed dish (MatTek, Ashland, MA, USA) covered with a thin sheet of 1% PB agar. Confocal fluorescence images were taken using an inverted confocal microscope (Nikon A1+) and Z-slices were obtained using a Ti Z-drive. See also SI Methods.

### Real-time PCR analysis

Dissociated slug cells were suspended in each solution, and shaken at 120 rpm, and harvested at the selected time points. Quantitative PCR amplification (qPCR) was performed using a qPCR thermocycler (ABI7500, Applied Biosystems) with primer pairs and fluorescent beacons (TaqMan probe MGB, Applied Biosystems) (Table S2). The levels of relative gene expression were calculated from the amplification value at the threshold cycle and the relative standard curve for each gene, followed by normalization by *rnlA* amplification as an endogenous control.

### Growth assay

Vegetative cells were starved at 2 × 10^6^ cells/cm^2^. *E. coli* suspension was applied either directly to the slugs or after mechanical dissociation. After 30h incubation, the samples were harvested and washed 2 times to remove the bacteria. Then, 3 × 10^6^ cells were transferred into 10 mL of the growth medium and shaken at 120 rpm. The cell number at different times was measured using a hemocytometer.

### Statistical analysis

*P*-values from multiple comparisons were adjusted using Holm’s method. A generalized linear model (GLM) and analysis of deviance for the fit were performed for the analysis listed in Table S1.

## Acknowledgments

We thank K. Kaneko for providing critical comments, T. Fujimori, T. Yoshida, and lab members for discussion and technical advice, and A. Nakajima for the microscopy setup. This work was supported by the Platform for Dynamic Approaches to Living System from the Ministry of Education, Culture, Sports, Science, and Technology, Japan to Y.S., M.S., and S.S.; and by a Grant-in-Aid for Challenging Exploratory Research (Nos. 16K14805) from the Japan Society for the Promotion of Science (JSPS) to Y.S., and M.S. and JSPS Grant-in-Aid 18H04759 and Japan Science and Technology Agency (JST) CREST JPMJCR1923 to S.S..

## Notes

### Competing Interest Statement

The authors have declared no competing interest.

### Summary of Updates

Minor changes to improve readability.

## References

[1] Hamilton WD. The genetical evolution of social behaviour. Journal of Theoretical Biology. 1964;7(1):1–52. doi:10.1016/0022-5193(64)90038-4

[2] Frank SA. Mutual policing and repression of competition in the evolution of cooperative groups. Nature. 1995;377(6549):520–522. doi:10.1038/377520a0

[3] West SA, Griffin AS, Gardner A, Diggle SP. Social evolution theory for microorganisms. Nature Reviews Microbiology. 2006;4(8):597–607. doi:10.1038/nrmicro1461

[4] Hallatschek O, Hersen P, Ramanathan S, Nelson DR. Genetic drift at expanding frontiers promotes gene segregation. Proceedings of the National Academy of Sciences of the United States of America. 2007;104(50):19926–19930. doi:10.1073/pnas.0710150104

[5] Buttery NJ, Jack CN, Adu-Oppong B, Snyder KT, Thompson CRL, Queller DC, Strassmann JE. Structured growth and genetic drift raise relatedness in the social amoeba *Dictyostelium discoideum*. Biology letters. 2012;8(5):794–797. doi:10.1098/rsbl.2012.0421

[6] Smukalla S, Caldara M, Pochet N, Beauvais A, Guadagnini S, Yan C, Vinces MD, Jansen A, Prevost MC, Latgé J-P, et al. *FLO1* is a variable green beard gene that drives biofilm-like cooperation in budding yeast. Cell. 2008;135(4):726–737. doi:10.1016/j.cell.2008.09.037

[7] Ho HI, Hirose S, Kuspa A, Shaulsky G. Kin recognition protects cooperators against cheaters. Current biology. 2013;23(16):1590–1595. doi:10.1016/j.cub.2013.06.049

[8] Velicer GJ, Kroos L, Lenski RE. Loss of social behaviors by *Myxococcus xanthus* during evolution in an unstructured habitat. Proceedings of the National Academy of Sciences of the United States of America. 1998;95(21):12376–12380.

[9] Danforth BN. Evolution of sociality in a primitively eusocial lineage of bees. Proceedings of the National Academy of Sciences. 2002;99(1):286–290. doi:10.1073/pnas.012387999

[10] James TY, Kauff F, Schoch CL, Matheny PB, Hofstetter V, Cox CJ, Celio G, Gueidan C, Fraker E, Miadlikowska J, et al. Reconstructing the early evolution of Fungi using a six-gene phylogeny. Nature. 2006;443(7113):818–822. doi:10.1038/nature05110

[11] Schirrmeister BE, Antonelli A, Bagheri HC. The origin of multicellularity in cyanobacteria. BMC Evolutionary Biology. 2011;11(1):45. doi:10.1186/1471-2148-11-45

[12] Wilson EO. The Insect Societies. Cambridge: Belknap Press; 1971. (Belknap Press).

[13] Wilson EO, Hölldobler B. Eusociality: Origin and consequences. Proceedings of the National Academy of Sciences of the United States of America. 2005;102(38):13367–13371. doi:10.1073/pnas.0505858102

[14] Libby E, Ratcliff WC. Ratcheting the evolution of multicellularity. Science. 2014;346(6208):426–427. doi:10.1126/science.1262053

[15] Libby E, Conlin PL, Kerr B, Ratcliff WC. Stabilizing multicellularity through ratcheting. Philosophical Transactions of the Royal Society B: Biological Sciences. 2016;371(1701):20150444. doi:10.1098/rstb.2015.0444

[16] Ratcliff WC, Denison RF, Borrello M, Travisano M. Experimental evolution of multicellularity. Proceedings of the National Academy of Sciences. 2012;109(5):1595–1600. doi:10.1073/pnas.1115323109

[17] Bull JJ, Charnov EL. On irreversible evolution. Evolution. 1985;39(5):1149–1155. doi:10.1111/j.1558-5646.1985.tb00455.x

[18] Santorelli LA, Thompson CRL, Villegas E, Svetz J, Dinh C, Parikh A, Sucgang R, Kuspa A, Strassmann JE, Queller DC, et al. Facultative cheater mutants reveal the genetic complexity of cooperation in social amoebae. Nature. 2008;451(7182):1107–1110. doi:10.1038/nature06558

[19] Khare A, Santorelli LA, Strassmann JE, Queller DC, Kuspa A, Shaulsky G. Cheater-resistance is not futile. Nature. 2009;461(7266):980–982. doi:10.1038/nature08472

[20] Manhes P, Velicer GJ. Experimental evolution of selfish policing in social bacteria. Proceedings of the National Academy of Sciences of the United States of America. 2011;108(20):8357–8362. doi:10.1073/pnas.1014695108

[21] Riley MA, Wertz JE. Bacteriocins: Evolution, Ecology, and Application. Annual Review of Microbiology. 2002;56(1):117–137. doi:10.1146/annurev.micro.56.012302.161024

[22] Danilova N. The evolution of immune mechanisms. Journal of Experimental Zoology Part B: Molecular and Developmental Evolution. 2006;306B(6):496–520. doi:10.1002/jez.b.21102

[23] Bonner JT. The Cellular Slime Molds. Princeton: Princeton University Press; 1967. doi:10.1515/9781400876884

[24] Kessin RH. Dictyostelium. Cambridge: Cambridge University Press; 2001. (Cambridge University Press). doi:10.1017/cbo9780511525315

[25] Strassmann JE, Zhu Y, Queller DC. Altruism and social cheating in the social amoeba *Dictyostelium discoideum*. Nature. 2000;408(6815):965–967. doi:10.1038/35050087

[26] Katoh M, Chen G, Roberge E, Shaulsky G, Kuspa A. Developmental Commitment in *Dictyostelium discoideum*. Eukaryotic Cell. 2007;6(11):2038–2045. doi:10.1128/ec.00223-07

[27] Raper KB. Pseudoplasmodium formation and organization in *Dictyostelium discoideum*. J Elisha Mitchell Sci Soc. 1940;56:241–282.

[28] Katoh M, Shaw C, Xu Q, Driessche NV, Morio T, Kuwayama H, Obara S, Urushihara H, Tanaka Y, Shaulsky G. An orderly retreat: Dedifferentiation is a regulated process. Proceedings of the National Academy of Sciences of the United States of America. 2004;101(18):7005–7010. doi:10.1073/pnas.0306983101

[29] Gilbert OM, Queller DC, Strassmann JE. Discovery of a large clonal patch of a social amoeba: implications for social evolution. Molecular Ecology. 2009;18(6):1273–1281. doi:10.1111/j.1365-294x.2009.04108.x

[30] Chen G, Zhuchenko O, Kuspa A. Immune-like Phagocyte Activity in the Social Amoeba. Science. 2007;317(5838):678–681. doi:10.1126/science.1143991

[31] Newell PC, Longlands M, Sussman M. Control of enzyme synthesis by cellular interaction during development of the cellular slime mold *Dictyostelium discoideum*. Journal of Molecular Biology. 1971;58(2):541–554.

[32] Deichsel H, Friedel S, Detterbeck A, Coyne C, Hamker U, MacWilliams HK. Green fluorescent proteins with short half-lives as reporters in *Dictyostelium discoideum*. Development Genes and Evolution. 1999;209(1):63–68.

[33] Siu CH, Roches BD, Lam TY. Involvement of a cell-surface glycoprotein in the cell-sorting process of *Dictyostelium discoideum*. Proceedings of the National Academy of Sciences of the United States of America. 1983;80:6596–6600.

[34] Siu CH, Harris TJC, Wang J, Wong E. Regulation of cell–cell adhesion during *Dictyostelium* development. Seminars in Cell & Developmental Biology. 2004;15(6):633–641. doi:10.1016/j.semcdb.2004.09.004

[35] Crowley TE, Nellen W, Gomer RH, Firtel RA. Phenocopy of discoidin I-minus mutants by antisense transformation in *Dictyostelium*. Cell. 1985;43(3):633–641. doi:10.1016/0092-8674(85)90235-1

[36] Bastounis E, Álvarez-González B, Álamo JC del, Lasheras JC, Firtel RA. Cooperative cell motility during tandem locomotion of amoeboid cells. Molecular Biology of the Cell. 2016;27(8):1262–1271. doi:10.1091/mbc.e15-12-0836

[37] Fujimori T, Nakajima A, Shimada N, Sawai S. Tissue self-organization based on collective cell migration by contact activation of locomotion and chemotaxis. Proceedings of the National Academy of Sciences of the United States of America. 2019;116(10):4291–4296. doi:10.1073/pnas.1815063116

[38] Steinberg MS, Takeichi M. Experimental specification of cell sorting, tissue spreading, and specific spatial patterning by quantitative differences in cadherin expression. Proceedings of the National Academy of Sciences of the United States of America. 1994;91(1):206–209. doi:10.1073/pnas.91.1.206

[39] Parkinson K, Bolourani P, Traynor D, Aldren NL, Kay RR, Weeks G, Thompson CRL. Regulation of Rap1 activity is required for differential adhesion, cell-type patterning and morphogenesis in *Dictyostelium*. Journal of Cell Science. 2009;122(3):335–344. doi:10.1242/jcs.036822

[40] Finney RE, Mitchell LH, Soll DR, Murray BA, Loomis WF. Loss and resynthesis of a developmentally regulated membrane protein (gp80) during dedifferentiation and redifferentiation in *Dictyostelium*. Developmental Biology. 1983;98(2):502–509. doi:10.1016/0012-1606(83)90379-2

[41] Lam TY, Pickering G, Geltosky J, Siu CH. Differential cell cohesiveness expressed by prespore and prestalk cells of *Dictyostelium discoideum*. Differentiation. 1981;20(1–3):22–28. doi:10.1111/j.1432-0436.1981.tb01151.x

[42] Whittingham WF, Raper KB. Non-viability of stalk cells in *Dictyostelium*. Proceedings of the National Academy of Sciences. 1960;46(5):642–649. doi:10.1073/pnas.46.5.642

[43] Harrington BJ, Raper KB. Use of a fluorescent brightener to demonstrate cellulose in the cellular slime molds. Applied Microbiology. 1968;16(1):106–113.

[44] Jermyn K, Traynor D, Williams J. The initiation of basal disc formation in *Dictyostelium discoideum* is an early event in culmination. Development. 1996;122:753–760.

[45] Watts DJ, Treffry TE. Culmination in the slime mould *Dictyostelium discoideum* studied with a scanning electron microscope. Journal of embryology and experimental morphology. 1976;35(2):323–333.

[46] Geritz SAH, Kisdi É, Meszéna G, Metz JAJ. Evolutionarily singular strategies and the adaptive growth and branching of the evolutionary tree. Evolutionary Ecology. 1998;12(1):35–57. doi:10.1023/a:1006554906681

[47] Dieckmann U, Ferrière R. Adaptive Dynamics and Evolving Biodiversity. In: Ferrière R, Dieckmann U, Couvet D, editors. Evolutionary conservation biology. Cambridge, UK: Cambridge University Press; 2004. p. 188–224.

[48] Smith JM. Evolution and the theory of games. Cambridge University Press; 1982. (Cambridge University Press).

[49] Eshel I. Evolutionary and continuous stability. Journal of Theoretical Biology. 1983;103(1):99–111. doi:10.1016/0022-5193(83)90201-1

[50] Sternfeld J. A study of PstB cells during *Dictyostelium* migration and culmination reveals a unidirectional cell type conversion process. Roux’s archives of developmental biology. 1992;201(6):354–363. doi:10.1007/bf00365123

[51] Abe T, Early A, Siegert F, Weijer C, Williams J. Patterns of cell movement within the *Dictyostelium* slug revealed by cell type-specific, surface labeling of living cells. Cell. 1994;77(5):687–699.

[52] Hochberg ME, Rankin DJ, Taborsky M. The coevolution of cooperation and dispersal in social groups and its implications for the emergence of multicellularity. BMC Evolutionary Biology. 2008;8(1):238. doi:10.1186/1471-2148-8-238

[53] Jack CN, Buttery N, Adu-Oppong B, Powers M, Thompson CRL, Queller DC, Strassmann JE. Migration in the social stage of *Dictyostelium discoideum* amoebae impacts competition. PeerJ. 2015;3:e1352. doi:10.7717/peerj.1352

[54] Nichols JM, Antolovic V, Reich JD, Brameyer S, Paschke P, Chubb JR. Cell and molecular transitions during efficient dedifferentiation. eLife. 2020;9:e55435. doi:10.7554/elife.55435

[55] Lande R. Expected time for random genetic drift of a population between stable phenotypic states. Proceedings of the National Academy of Sciences of the United States of America. 1985;82(22):7641–7645.

[56] Iwasa Y, Michor F, Nowak MA. Stochastic tunnels in evolutionary dynamics. Genetics. 2004;166(3):1571–1579.

[57] Sternfeld J. The anterior-like cells in *Dictyostelium* are required for the elevation of the spores during culmination. Development Genes and Evolution. 1998;208(9):487–494.

[58] Sternfeld J, David CN. Cell Sorting daring Pattern Formation in *Dictyostelium*. Differentiation. 1981;20(1–3):10–21. doi:10.1111/j.1432-0436.1981.tb01150.x

[59] Dormann D, Siegert F, Weijer CJ. Analysis of cell movement during the culmination phase of *Dictyostelium* development. Development. 1996;122(3):761–769.

[60] Saito T, Kato A, Kay RR. DIF-1 induces the basal disc of the *Dictyostelium* fruiting body. Developmental Biology. 2008;317(2):444–453. doi:10.1016/j.ydbio.2008.02.036

[61] Bichler G, Weijer CJ. A *Dictyostelium* anterior-like cell mutant reveals sequential steps in the prespore prestalk differentiation pathway. Development. 1994;120(10):2857–2868.

[62] Smith JM, Szathmáry E. The Major Transitions in Evolution. New York: Oxford University Press; 1995. (Oxford University Press).

[63] Michod RE, Roze D. Cooperation and conflict in the evolution of individuality. III. Transitions in the unit of fitness. In: Nehaniv CL, editor. Mathematical and Computational Biology: Computational Morphogenesis, Hierarchical Complexity, and Digital Evolution. American Mathematical Society; 1999. p. 47–92.

[64] Hauert C, Monte SD, Hofbauer J, Sigmund K. Replicator dynamics for optional public good games. Journal of Theoretical Biology. 2002;218:187–194. doi:10.1006/yjtbi.3067

[65] Garcia T, Doulcier G, Monte SD. The evolution of adhesiveness as a social adaptation. eLife. 2015;4:e08595. doi:10.7554/elife.08595.001

[66] Dubravcic D, Baalen M van, Nizak C. An evolutionarily significant unicellular strategy in response to starvation in *Dictyostelium* social amoebae. F1000Research. 2014;3:133. doi:10.12688/f1000research.4218.2

[67] Tarnita CE, Washburne A, Martinez-Garcia R, Sgro AE, Levin SA. Fitness tradeoffs between spores and nonaggregating cells can explain the coexistence of diverse genotypes in cellular slime molds. Proceedings of the National Academy of Sciences of the United States of America. 2015;112(9):2776–2781. doi:10.1073/pnas.1424242112

[68] Kuzdzal-Fick JJ, Foster KR, Queller DC, Strassmann JE. Exploiting new terrain: an advantage to sociality in the slime mold *Dictyostelium discoideum*. Behavioral Ecology. 2007;18(2):433–437. doi:10.1093/beheco/arl102

