## Supplementary Information Text for "Evolutionary stability of social commitment"

### Supplementary Information for
### Evolutionary stability of social commitment

Yuka Shirokawa*, Masakazu Shimada, Nao Shimada, Satoshi Sawai

*To whom correspondence should be addressed: Yuka Shirokawa


**This PDF file includes:**

Supplementary text
Figures S1 to S7
Tables S1 to S3
Legends for Movies S1 to S5
Legends for Datasets S1 and Code S1
SI References

**Other supplementary materials for this manuscript include the following:**

Movies S1 to S5
Datasets S1
Code S1

**Supplementary Information Text**

**Strain construction**

Cells were grown at 22°C in a shaking culture with PS growth medium [1]. To obtain cells that co-expressed prespore and prestalk fate markers, the following two constructs were introduced into AX4 cells, and clones were isolated: A construct with G418 selection that confers RFPmars expression under the control of the prestalk-specific *ecmAO* promoter (ecmAO-RFP, Dicty Stock Center, plasmid ID 639): A GFP expression construct under the control of the prespore-specific *pspA* promoter with hygromycin selection (pspA-GFP). For selection and plasmid maintenance, 30 µg/mL G418 and 60 µg/mL hygromycin were added to the growth media. The pspA-GFP expression plasmid was constructed by replacing the *act15* promoter of pHygGFP [2] with the *pspA* promoter as follows: A *Cla*I restriction site in pHygGFP was replaced with a *Bgl*II site using the oligonucleotide linker 5'-CGACAGATCTGT-3'. The *act15* promoter was then replaced with the *pspA* promoter at *Xba*I and *Bgl*II sites in the pHygGFP. The *pspA* promoter fragment was obtained from the pspA-Gal construct (Dicty Stock Center plasmid ID 49) [3] by excision at *Xba*I and *Bgl*II sites. Strains constitutively expressing GFP or RFP were constructed by transforming AX4 with GFP or RFP expression constructs under the control of the *act15* promoter, with 10 µg/mL G418 selection. For GFP expression, pA15GFP (S65T) [4] was used. For RFP expression, pA15-mRFPmars [5] was used.

**Refeeding experiments**

Vegetative cells were washed with phosphate buffer (PB) (pH 6.5; 20 mM $KH_2PO_4$, 20 mM $Na_2HPO_4$) and plated on a 1% PB agar (BactoAgar, Difco) plate at 0.3 to 0.4 $\times 10^6$ cells/$cm^2$ and incubated at 22°C for 18 h until the slug stage. Cells were collected by mechanical dissociation in PB containing 20 mM EDTA (pH 6.4) by repeated pipetting and passage through a 23 G needle and a 40µm cell strainer followed by washing with PB.

To prepare a bacterial suspension, *E. coli* B/r was grown in LB medium (10 g/L tryptone, 5 g/L yeast extract, and 10 g/L NaCl) to the optical density (OD) of 3 at 600 nm, and was concentrated in PB at OD50 equivalent by centrifugation. The bacterial suspension was used immediately. For refeeding, dissociated cells were co-suspended at 2 to 3 $\times 10^6$ cells/ml final density with *E. coli* B/r at OD50 in PB and shaken for 3 or 5 h in a 50 ml centrifuge tube. The typical volume was 1.25 mL. The cells were then washed by pelleting and resuspending in PB; this procedure was repeated 2 times to remove bacteria. In the case of refeeding with growth media, the PS growth medium was used. NF cells were cells suspended in PB without nutrients immediately after dissociation.

For mixing experiments (Figure 1,2), NF and RF cells were mixed at 85:15 ratio. The mixed cell suspension adjusted to 1 to 2 $\times 10^6$ cells/mL was plated on a glass-bottomed dish (MatTek, Ashland, MA) covered with a thin sheet of 1% PB agar. The plates were left still for 15 min to allow the cells to attach to the agar before the buffer was gently removed. Confocal images of GFP, RFP, and calcofluor white fluorescence were taken using an inverted confocal microscope (A1+, Nikon) and Z-slice was obtained using Ti Z-drive. The cells were observed at three-time windows after plating: 1.5–2 h for the aggregates with 20× air objective lens, 4–5 h during early culmination for the upper part of a fruiting body with 40× air objective lens, 8–10 h for the basal disk of terminally differentiated fruiting bodies with 60× oil immersion objective lens, and for cells scattered around the base of fruiting bodies with 20× air objective lens.

Images were analyzed using custom programs in ImageJ [6] and the R language [7]. To determine and quantify cell localization, images were binarized to obtain masks according to the fluorescence of cell fate markers. For cell aggregates, lists of XY coordinates of marker-positive pixels were obtained from the binarized images. Z-sections at 3 µm intervals taken between 30 to 60 µm from the bottom of the aggregate were analyzed. Lists of XY coordinates from each slice were merged into a single list. The origin of the coordinate was set to the aggregate center and the aggregate radius was normalized to 1. The average distance from the center to marker-positive pixels was computed for the prestalk marker and prespore marker.

To capture images of early culminant, the upper regions of fruiting bodies were attached to a glass slide and Z-slices were acquired at 5 µm intervals from the bottom to the top of a sample. Lists of XY coordinates of marker-positive pixels from Z-slices were merged. Each fruiting body was aligned vertically from the tip to the lower cup, and a normalized coordinate in the vertical direction was assigned from 0 to 1. The frequency of positive pixels at each position was quantified for the prestalk and prespore markers. The vertical position was divided into 50 equally-spaced bins to obtain histograms.

For the analysis of the basal disk, stalks were transferred to a glass slide and stained with 0.01% calcofluor white (BD, NJ, USA). Basal disk regions were determined by calcofluor white fluorescence which stains cellulose from vacuolated stalk cells [8]. Z-section images were acquired at a 1.5µm interval from the bottom to the top of the sample. The number of marker-positive pixels in the region was acquired from binarized images and the values from each slice were summed. The ratio of the pixel number between the prestalk and prespore images was calculated.

For cells scattered around the base of fruiting bodies, the standing fruiting body on an agar-coated glass-bottom dish was observed from the bottom of the dish. Z-stack images were collected from the bottom of the sample to a height of 30 µm at 5 µm intervals. The number of marker-positive pixels was acquired within a 100 µm radius from the basal disk center, and the values from each Z-slice were summed. Ratios of the values between the prestalk and prespore images were calculated.

**Quantitative real-time polymerase chain reaction analysis**

qRT-PCR analysis was performed as follows. For ‘No-dissociation’ sample (Figure 1*D*), AX4 vegetative cells were starved on a 1% PB agar plate and harvested at the selected time points. For the other conditions (‘No-nutrient buffer’, ‘Bacteria’, ‘Growth medium’, in Figure 1*D*), dissociated slug cells were suspended in PB or PB together with *E. coli* at OD50 or the PS growth medium and shaken in a 50 ml centrifuge tube. Cells were harvested at the selected time points, and total RNA was extracted using a Maxwell 16 LEV simplyRNA cells and Tissue kit (Promega, WI, USA). cDNA was synthesized using random hexamers and SuperScript III (First-Strand Synthesis, Invitrogen). Quantitative PCR (qPCR) amplification was performed with a qPCR thermocycler (ABI7500, Applied Biosystems) using a pre-mixed reaction solution (TaqMan Universal PCR Master Mix, Applied Biosystems) and with primer pairs and fluorescent beacons (TaqMan probe MGB, Applied Biosystems) (Table S2). The amplification value at the threshold cycle (CT) of each sample was measured in three independent wells, and the average was used. The levels of relative gene expression were calculated from the CT and the relative standard curve for each gene followed by normalization by *rnlA* amplification as an endogenous control.

For hierarchical clustering of time series of the gene expression level, permutation distribution clustering was applied with R function ‘pdc’. Time series of the three conditions (‘No-nutrient buffer’, ‘Bacteria’, and ‘Growth medium’) were embedded and calculated the squared Hellinger distance to measure dissimilarity between gene targets, and then performed clustering according to the dissimilarity.

To investigate temporal differences in the level of gene expression, ‘hypothesis testing with bootstrap’ [9] was applied to the qRT-PCR data. Pair-wise comparisons were performed between the sampling time points 0h and 5h. The null hypothesis was that the two samples originated from the same probability distribution. Pairs of 100,000 bootstrap samples were generated from a mixture of the observed data. The percentile rank of the observed *t*-values within the distribution of *t*-values was generated from the bootstrap sampling. And then, the percentile rank was used as the *p*-value for a two-sided test.

**Cell cohesiveness assay**

Cell cohesiveness was assayed as described [10] with minor modifications in the cell density and the container geometry. Dissociated slugs with the cell fate markers were suspended at $6 \times 10^6$ cells/mL in plain PB or PB with *E. coli* at OD50 or the PS growth medium, placed in 50 mL tubes and shaken at 120 rpm for 1.5, 3, 5 h. Since attachment of bacteria to *Dictyostelium* cells can inhibit cohesion between *Dictyostelium* cells, refeeding in the liquid growth medium was also tested for refeeding conditions. For RF conditions, the assay was performed under the continued presence of nutrients to avoid the chemotaxis effect by cAMP. The number of cells that were not associated with aggregates was quantified as follows: Images of the sample loaded in a hemocytometer were acquired and binarized according to the fluorescence of the cell fate markers. The number of single cells was then calculated using the particle analyzer function in ImageJ.

**Flow cytometry**

Dissociated cells with the cell fate markers were suspended in PB together with *E. coli* B/r at OD50 and shaken for 3, 6, and 12 h. The cells were washed with PB and the intensity of RFP and GFP expression was measured using a flow cytometer (SH800, Sony, Japan). “Prespore fate” denotes negative for ecmAO-RFP fluorescence and positive for pspA-GFP fluorescence. “Prestalk fate” is positive for ecmAO-RFP fluorescence and negative for pspA-GFP fluorescence. “Non-fluorescent cells” denotes negative for ecmAO-RFP fluorescence and negative for pspA-GFP fluorescence.

**Terminal cell fate allocation in the mixed cell population**

Allocation to sporulation and solitary amoeboid cells of RF cells in the terminal state was quantified. We mixed at 1:1 ratio NF and RF cells constitutively expressing GFP or RFP under the strong actin15 promoter, respectively. The cell mixtures were applied on a 1% PB agar plate. After incubation at 22°C for 10 h, when development of the fruiting body has more or less completed, PB containing 20 mM EDTA was poured onto the agar plate and non-vacuolated cells i.e., spores and solitary amoeboid cells were harvested by pipetting. The cell suspension was treated with 0.01% calcofluor white (BD, NJ, USA) to distinguish spores from amoeboid cells [8]. The number of spores and amoeboid cells was counted, and that of refed and non-refed cells was identified according to GFP and RFP expression. The GFP-RFP ratio in the terminal state was normalized by the GFP-RFP ratio of dissociated cells immediately after mixing. The results were shown in Figure S4.

**Growth assay to measure growth cost of the social commitment**

Vegetative cells were washed with PB to remove nutrients and plated at $2 \times 10^6$ cells/cm$^2$ on a Whatman no. 50 filter on four sheets of Whatman no. 3 filter soaked with PB and incubated at 22°C. After 18h, *E. coli* B/r at OD400 in PB was applied directly to developing slugs on the plates or dissociated slugs. The dissociated cells were replated on new filters. After 48 h from the initial time of starvation (30 h from the slug stage), cells (including amoeboid cells and spores) were harvested by shaking the filter with PB in 50 mL tubes. The cells were then washed 2 times to remove bacteria. For the growth assay, $3 \times 10^6$ cells were transferred to 10 ml PS growth medium, and incubated in a shaken culture at 120 rpm. The number of cells was counted using a hemocytometer over time.

To estimate the growth delay incurred by fruiting body formation ($d$, Figure 3*A*), the number of amoeboid cells immediately after the cell harvesting (30 h after from the slug stage) is compared among conditions. To estimate the lag time for spore germination ($\lambda$, Figure 3*A*), the cell growth within the growth medium was used. Growth curves were fitted with the following logistic model [11],

$$y = \frac{D}{(1+\exp\left(\frac{4j}{D}(lag-t)+2\right))},$$

where $y$ is the cell number, $D$ is the maximum cell number, and $t$ is the time, $lag$ is the lag time between the cell inoculation and initiation of cell growth, $j$ is the growth rate. The parameters $j$, $lag$, $D$ were estimated using nonlinear least squares function 'nls' in R. The results were shown in Table S1.

**Statistical analysis**

All statistical analyses were performed in R [7]. Details of a generalized linear model (GLM) and analysis of deviance for the fit were listed in Table S1. For multiple comparisons, the *p*-value was adjusted with Holm's method.

### Evolutionary model

For valuables and parameters, see Table S3. All numerical calculations for plotting of the model were performed with Mathematica.

**Evolutionary transition of mutant cell fate**

We consider three types of social systems (canonical, reverse, and random allocation systems). In the canonical allocation system, the evolutionary changes of the mutant cell fate can be described by the differential equation Eq. **1** in the main text.

In the reverse allocation systems, the mutant cell fate transitions from prestalk, to prespore, and then solitary fate via the evolution of $x$. Thus, the evolutionary changes can be described by swapping the variables in Eq. **1** so that

$$\frac{dP(SP|x)}{dx} = -k_1 P(ST|x) + k_2 P(SP|x),$$
$$\frac{dP(ST|x)}{dx} = k_1 P(ST|x), \qquad \text{(S1)}$$
$$\frac{dP(Sol|x)}{dx} = -k_2 P(SP|x),$$

where, $k_1$, $k_2$ are the rates of cell fate transition ($k_1 > 0$, $k_2 > 0$). We assume $k_1 \gg k_2$ in the reverse allocation systems, where the default values are $k_1 = 23.3$, $k_2 = 4$ (see also, SI text, 'Default $k1$, $k2$ values'). The equation Eq. **S1** is solved using the initial value at $x = 1$ (prespore: $SP_{initial} = 0.6$, prestalk: $ST_{initial} = 0.2$, solitary: $Sol_{initial} = 0.2$ ).

In the random allocation system, the mutant cell fate randomly transitions to a solitary cell from a prestalk or prespore cell. Therefore, the evolutionary changes are,

$$\frac{dP(SP|x)}{dx} = k_1 P(SP|x),$$
$$\frac{dP(ST|x)}{dx} = k_2 P(ST|x), \qquad \text{(S2)}$$
$$\frac{dP(Sol|x)}{dx} = -k_1\, P(SP|x) - k_2 P(ST|x),$$

In the random allocation system, prespore and prestalk cells exhibit the same degree of cohesion (prespore = prestalk > solitary cell), therefore a prespore and prestalk cell are assumed to have the same transition rate $k_1 = k_2$. We set the default values as $k_1 = 3.9, k_2 = 3.9$.

**Selection gradient and evolutionary singular point**

Invasion fitness in Eq. **2** in the main text is differentiable, and the selection gradient is

$$\left.\frac{\partial W(x';x_{wt})}{\partial x'}\right|_{x'=x} = \{(\frac{\partial P(SP|x')}{\partial x'} + g\frac{\partial P(Sol|x')}{\partial x'}) - (\frac{\partial P(SP|x_{wt})}{\partial x'} + g\frac{\partial P(Sol|x_{wt})}{\partial x'})\}|_{x'=x}. \quad \text{(S3)}$$

The point where the selection gradient vanishes, i.e. "evolutionarily singular" [12, 13] is denoted as $x^*$, and calculated from Eq. **S3** when this equation is set to zero:

$$\left.\frac{\partial W(x';x_{wt})}{\partial x'}\right|_{x'=x} = 0. \quad \text{(S4)}$$

The singular point was described as the following format,

$$x^* = \frac{Log(\frac{B}{A})}{k_1 - k_2}, \quad \text{(S5)}$$

In the canonical allocation system, from Eq. **S4** and Eq. **1** in the main text,

$$A = \frac{k_1 k_2 SP_{initial}(k_1 - k_2 + k_2 g)e^{-k_1}}{k_1 - k_2},$$

$$B = \frac{-(k_2)^2 g(-k_1 SP_{initial} - k_1 ST_{initial} + k_2 ST_{initial})e^{-k_2}}{k_1 - k_2}.$$

In the reverse allocation system, from Eq. **S4** and Eq. **S1**,

$$A = \frac{k_1 k_2 ST_{initial}(k_2 g - k_1)e^{-k_1}}{k_1 - k_2},$$

$$B = \frac{(k_2)^2(1-g)(-k_1 SP_{initial} + k_2 SP_{initial} - k_1 ST_{initial})e^{-k_2}}{k_1 - k_2}.$$

In the random allocation system, from Eq. **S4** and Eq. **S2**,

$$A = k_1 SP_{initial}(1-g)e^{-k_1},$$

$$B = k_2 g ST_{initial} e^{-k_2},$$

The analytical predictions of the singular point are plotted in Figure 4*D*.

**Condition for ESS and convergence stable**

The condition for ESS [14] of the singular strategy $x^*$ is

$$\left.\frac{\partial^2 W(x';x_{wt})}{\partial x'^2}\right|_{x=x'=x^*} < 0. \quad \text{(S6)}$$

The criteria for convergence stable [12, 15] of the singular strategy $x^*$ is

$$\left.\frac{\partial^2 W(x';x_{wt})}{\partial x^2}\right|_{x=x'=x^*} - \left.\frac{\partial^2 W(x';x_{wt})}{\partial x'^2}\right|_{x=x'=x^*} > 0. \quad \text{(S7)}$$

In the canonical allocation system,

$$\left.\frac{\partial^2 W(x';x_{wt})}{\partial x'^2}\right|_{x=x'=x^*} = P(SP|x^*)k_1(k_1 - k_2 + k_2 g), \quad \text{(S8)}$$

$$\left.\frac{\partial^2 W(x';x_{wt})}{\partial x^2}\right|_{x=x'=x^*} - \left.\frac{\partial^2 W(x';x_{wt})}{\partial x'^2}\right|_{x=x'=x^*} = -2P(SP|x^*)k_1(k_1 - k_2 + k_2 g). \quad \text{(S9)}$$

Therefore, if

$$k_1 < k_2(1-g), \quad \text{(S10)}$$

both Eq. **S6** and Eq. **S7** are satisfied. However, Eq. **S10** is never satisfied because we assume $k_1 \gg k_2$. This means that Eq. **S8** is always positive, and the singular point acts as an evolutionary repellor and not ESS. In addition, Eq. **S9** is always negative; therefore, the singular point is not convergence stable. The numerical calculations changing $k_1$, $k_2$ and $g$ (Figure S7) shows consistent results.

In the reverse allocation system,

$$\left.\frac{\partial^2 W(x';x_{wt})}{\partial x'^2}\right|_{x=x'=x^*} = P(SP|x^*)(1-g)k_2(k_2 g - k_1), \tag{S11}$$

$$\left.\frac{\partial^2 W(x';x_{wt})}{\partial x^2}\right|_{x=x'=x^*} - \left.\frac{\partial^2 W(x';x_{wt})}{\partial x'^2}\right|_{x=x'=x^*} = -2P(SP|x^*)(1-g)k_2(k_2 g - k_1). \tag{S12}$$

Therefore, under either of the following conditions,

$$g < 1 \text{ and } k_2 g < k_1, \tag{S13}$$

$$g > 1 \text{ and } k_2 g > k_1, \tag{S14}$$

both Eq. **S6** and Eq. **S7** are satisfied. Under starved conditions $g < 1$, Eq. **S13** is always satisfied because we assume $k_1 \gg k_2$. This means that Eq. **S11** is negative, and the singular point is an attractor and ESS. In addition, Eq. **S12** is always positive; therefore, the singular point is convergence stable. On the other hand, under nutrient-rich conditions $g > 1$, there is no singular point in $0 \leq x \leq 1$ (Figure S7).

In the random allocation system, there is no singular point under the assumed condition ($k_1 = k_2$)(Eq. **S5**).

**An alternative formulation of the invasion fitness as explicit altruism.**

In the main text, invasion fitness is expressed as a result of resource allocation between solitary and social fitness (Eq. **3**). On the other hand, social fitness can also be expressed as a result of altruism, where successful sporulation depends on the average stalk investment in a group (a fruiting body). For the wild-type cell, the average stalk investment is simply $P(ST|x_{wt})$, and for the rare mutant cell in a group of size $n$, that is $(P(ST|x') + (n-1)P(ST|x_{wt}))/n$. Therefore, the fitness of the wild-type cell is,

$$f(x_{wt};x_{wt}) = P(SP|x_{wt})P(ST|x_{wt}) + g\,P(Sol|x_{wt}), \tag{S15}$$

the mutant fitness is,

$$f(x';x_{wt}) = P(SP|x')\left(\frac{P(ST|x') + (n-1)P(ST|x_{wt})}{n}\right) + g\,P(Sol|x'). \tag{S16}$$

The result (Figure S6) is fundamentally the same as that in Figure 4*C*, for the fitness plot reveals a fitness valley. The valley is slightly shallower ($g = 0.3, 1$) than depicted in Figure 4, and this is likely because stalk differentiation contributes to spore success, fitness load of the stalk is reduced.

**Suitability of default cell fate ratio**

We set the default cell fate values as $SP_{initial} = 0.6$, $ST_{initial} = 0.2$, and $Sol_{initial} = 0.2$, respectively. Although the above analysis indicates that our conclusion does not depend on the specific choice in the cell-type ratio, the flow cytometric analysis (Figure S5) supported our choice of the default parameter set. We detect 60% of prespore cells and 20% of prestalk cells in the cell population of the slug stage. Besides these cells, we also found 20% of non-fluorescent cells in the population. We consider that most of these cells are solitary cells that are known not to differentiate under starvation, such as 'non-aggregating cells' [16], and cells leaves behind in the tail of slugs [17]. The non-aggregating cells are estimated as 15-20% of the total population [16].

**Default $\boldsymbol{k1}$, $\boldsymbol{k2}$ values**

For biological reality, we set default $k_1$, $k_2$ values that mimic the cell fate transition patterns associated with a change in a specific cohesion distribution as the conceptual illustration in Figure 4*A*. Briefly, we consider normal distributions of cohesion $c$ for prespore ($sp$), prestalk ($st$), and solitary cells ($sol$). The evolution of $x$ is assumed to decrease the means of the cohesion distributions. Based on the original distribution of a wild-type cell ($x = 1$, full social commission), inflow (i.e., the increment of solitary fitness due to the evolution of $x$) and outflow (i.e., the decrement of social fitness due to the evolution of $x$) were calculated from the distribution shifts. We determined appropriate $k_1$ and $k_2$ values by making an equation that describes the equality between the cell fate probability change due to the distribution shifts and that in the differential equations of Eq. **1**. And then, we solved this with respect to $k_1$ and $k_2$. These solutions then allowed us to set the default values as follows: $k_1 = 14.6$, $k_2 = 6.3$ (the canonical allocation system), $k_1 = 23.3$, $k_2 = 4$ (the reverse allocation system), $k_1 = 3.9$, $k_2 = 3.9$ (the random allocation system). Again, the conditions for ESS and convergence stable is not affected by the default $k_1$, $k_2$ under our assumed condition ($k_1 \gg k_2$). See also Figure S7.

**Cheating suppression through slug migration**

An evolutionary trait $i$ is additional spore investment at the slug stage, thus the initial cell fate probabilities for prespore, prestalk, and solitary cells, are $SP_{initial} + i$, $ST_{initial} - i$, and $Sol_{initial}$, respectively.

Cell fate transition over time through the slug migration can be described as a series of differential equations with derivative with respect to time $t$.

In the canonical allocation system, the equation is

$$\frac{dP_t(SP)}{dt} = -k_1 P_t(SP),$$

$$\frac{dP_t(ST)}{dt} = k_1 P_t(SP) - k_2 P_t(ST), \qquad \text{(S17)}$$

$$\frac{dP_t(Sol)}{dt} = k_2 P_t(ST),$$

where, $P_t(Z)$ is the probability of taking the fate $Z$ at the time $t$ (SP: Spore, ST: Stalk, Sol: Solitary cell). Both the wild-type and cheating mutant cell follow Eq. **S17**.

In the reverse allocation system, the cell fate transition is

$$\frac{dP_t(SP)}{dt} = k_1 P_t(ST) - k_2 P_t(SP),$$

$$\frac{dP_t(ST)}{dt} = -k_1 P_t(ST), \qquad \text{(S18)}$$

$$\frac{dP_t(Sol)}{dt} = k_2 P_t(SP).$$

In the random allocation system, the equation is

$$\frac{dP_t(SP)}{dt} = -k_1 P_t(SP),$$

$$\frac{dP_t(ST)}{dt} = -k_2 P_t(ST), \qquad \text{(S19)}$$

$$\frac{dP_t(Sol)}{dt} = k_1 P_t(SP) + k_2 P_t(ST).$$

The differential equations (Eq. **S17**, Eq. **S18**, and Eq. **S19**) were solved with respect to $t$ using initial conditions at $t = 0$ (prespore: $P_0(SP) = SP_{initial} + i$, prestalk: $P_0(ST) = ST_{initial} - i$, solitary: $P_0(Sol) = Sol_{initial}$). The solution is designated as $P_t(SP|i)$, $P_t(ST|i)$, and $P_t(Sol|i)$ respectively.

Based on previous experimental studies [17, 18], cell fate transition via the migration is assumed to maintain a constant cell fate proportion, thus a loss of a prestalk cell from the slug tail is compensated for by trans-differentiation of a prespore to a prestalk cell. For the constant proportioning, the ratio between $k_1$ and $k_2$ ($k_2 = \omega\, k_1$) is obtained as satisfying the following condition,

$$\frac{P_t(SP|i=0)}{P_t(ST|i=0)} = \beta, \qquad \text{(S20)}$$

where $\beta$ is a constant with respect to $t$.

In the canonical allocation system,

$$\omega = \frac{SP_{initial} + ST_{initial}}{ST_{initial}}.$$

In the reverse allocation system,

$$\omega = \frac{SP_{initial} + ST_{initial}}{SP_{initial}}.$$

In the random allocation system, $\omega = 1$.

Based on the default value ($SP_{initial} = 0.6$, $ST_{initial} = 0.2$, and $Sol_{initial} = 0.2$), we set the value as $k_1 = 6$, $k_2 = 24$ in the canonical allocation system, $k_1 = 4.5$, $k_2 = 6$ in the reverse allocation system, $k_1 = 6$, $k_2 = 6$ in the random allocation system.

The invasion fitness of the cheating mutant cell is,

$$W(i'; i_{wt}) = f(i'; i_{wt}) - f(i_{wt}; i_{wt}), \quad \text{(S21)}$$

where, $f(i'; i_{wt})$ is the survival probability of the mutant cell, and $f(i_{wt}; i_{wt})$ is that of the wild-type cell. Both $f(i'; i_{wt})$ and $f(i_{wt}; i_{wt})$ are defined as the sum of the cell fate probabilities as follows,

$$f(i) = P_t(SP|i) + g\, P_t(Sol|i). \quad \text{(S22)}$$

Eq.**22** indicates that the survival probability is defined based on its own trait value, without complex chimeric interactions.

The selection gradient is

$$\left.\frac{\partial W(i'; i_{wt})}{\partial i'}\right|_{i'=i}. \quad \text{(S23)}$$

The selection gradient was always constant with respect to $i$, irrespective of the allocation system. Thus, if the selection gradient is positive, the cheating mutant can evolve and acquire maximum spore investment $i$.

In the canonical allocation system, the cheating mutant evolves if

$$g < \frac{e^{k_2 t}(k_2 - k_1)}{k_2(e^{k_2 t} - e^{k_1 t})}. \quad \text{(S24)}$$

In the reverse allocation system, the mutant evolves if

$$g > \frac{(k_2 e^{k_1 t} - k_1 e^{k_2 t})}{k_2(-e^{k_2 t} + e^{k_1 t})}. \quad \text{(S25)}$$

On the other hand, in the random allocation system, under assumed condition ($k_2 = k_1$),

$$\left.\frac{\partial W(i'; i_{wt})}{\partial i'}\right|_{i'=i} = e^{-k_1 t}. \quad \text{(S26)}$$

Therefore, the selection gradient is always positive in the random allocation system. The analytical predictions for cheater evolution are plotted in Figure 5.

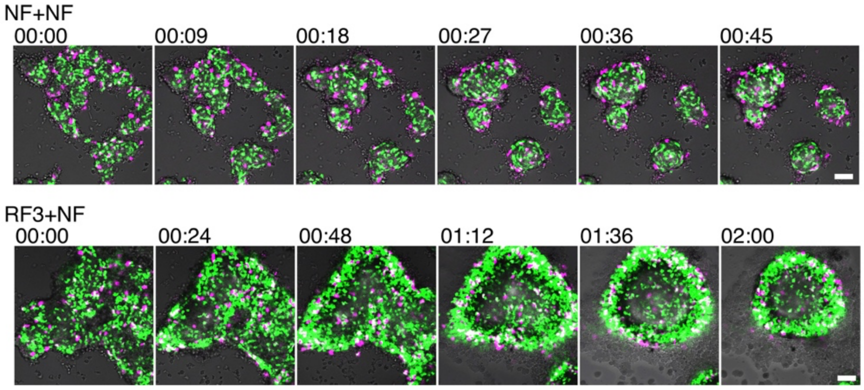


**Fig. S1. The time course of the reaggregation of dissociated cells.**

Time series taken 1.5–2 h after plating. The merged fluorescent and bright field images. In NF+NF, prestalk cells (ecmAO-RFP, magenta) were sorted to the peripheral region of the aggregate, while prespore cells (pspA-GFP, green) were more uniformly distributed. For RF3+NF, RF cells of both prestalk and prespore origins were well mixed and then sorted out to the periphery of the aggregates. The numbers on the upper left side indicate hour:min. All scale bars are 50μm.

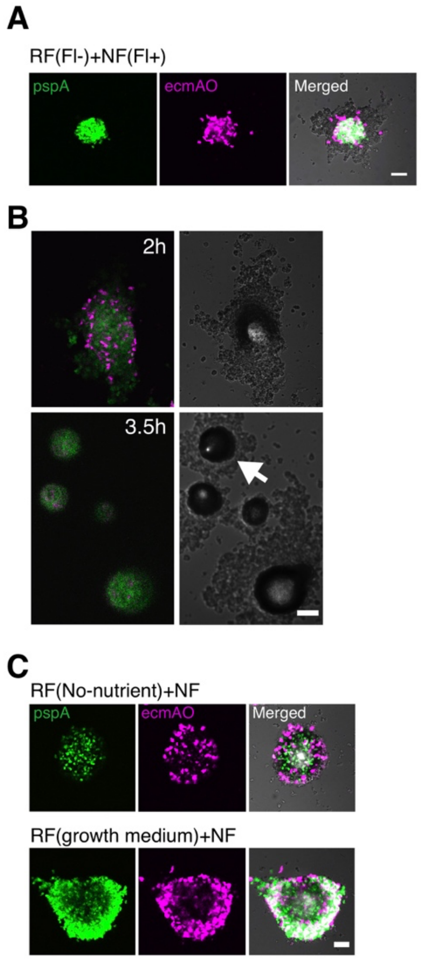


**Fig. S2. Supporting data for refeeding experiments.**

(*A*) The swap control: NF cells with the cell-type reporter (fluorescent) and nonfluorescent RF3 cells were mixed at a ratio 15:85 (RF(Fl-)+NF(Fl+)). Representative images of mixed aggregates (pspA: GFP-channel. ecmAO: RFP-channel. Merged: overlay of bright-field (grayscale) and fluorescence images). A reciprocal pattern of fluorescent cells in the center and non-fluorescent cells in the periphery were shown. The result indicates that expression of the marker genes itself did not affect the sorting pattern.

(*B*) RF3 cells were plated alone and observed aggregation. A merged fluorescent image of the prespore (green) and prestalk (magenta) marker expression (left), and bright field images (right). Upper panel: 2h after plating. Lower panel: 3.5h after plating. RF cells formed a mound and tip (arrow) at roughly the same time as NF cells. The result indicate that RF cells were still fully capable of developing on their own, suggesting that segregation was not due to the innate inability of RF cells to develop, but rather a relative behavior when in association with the NF cells.

(*C*) 'RF(No-nutrient)+NF' and 'RF(growth medium)+NF': Cells (fluorescent) shaken for 3 h in the plain phosphate buffer or the growth medium were mixed with NF cells (nonfluorescent). RF(No-nutrient)+NF show that cells dissociated and shaken without nutrients did not segregate from the freshly dissociated cells. RF(growth medium)+NF indicates that refeeding with liquid growth medium had the same effect as refeeding with bacteria. These results indicates that the unique sorting pattern was not merely due to mechanical interruption but also required nutrient replenishment. All scale bars = 50 µm.

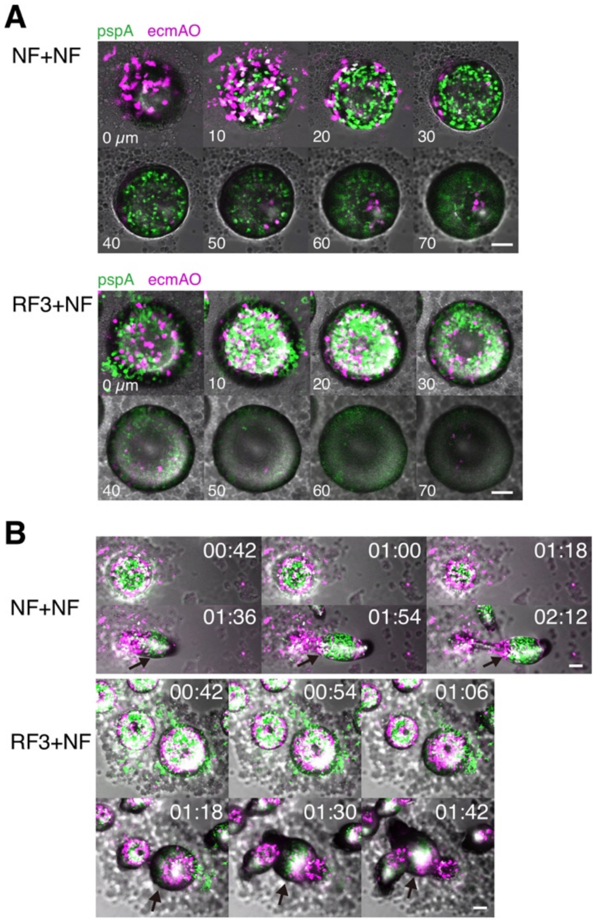


**Fig. S3. Fruiting body formation of refed cells.**

(*A*) Z stack images of an aggregate in the tip formation stage (3h after plating). Upper panels: NF+NF. Lower panels: RF3+NF. The numbers in the images are the height from the bottom of an aggregate (µm). Under NF+NF condition, prestalk cells localized to the bottom of the aggregate and prespore cells to the upper regions. In RF3+NF condition, RF cells of both prestalk and prespore origins localized to the bottom.

(*B*) The time series of fruiting body formation from 4-5h after plating (maximum intensity projections). Note that in NF+NF, prespore cells occupied the middle region (arrows). In the RF3+NF, refed cells localized to the lower cup position of the fruiting body. These outcomes indicate that refed prespore cells lost their ability to occupy the upper position of the fruiting body. The numbers on the upper right side are hour:min. All scale bars are 50µm.

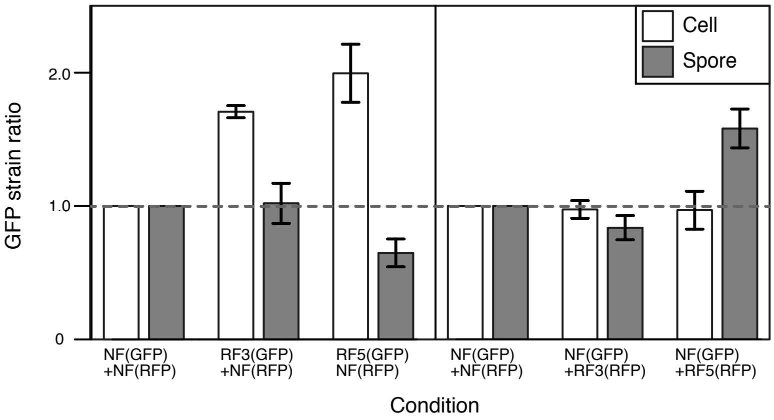


**Fig. S4. Terminal cell fate allocation in the mixed cell population.**

Non-refed and refed cells constitutively express GFP (act15-GFP) or RFP (act15-RFP).

Left panel: The mixtures of refed GFP cells with non-refed RFP cells (RF3(GFP)/NF(RFP), RF5(GFP)/NF(RFP)). Right panel: The mixtures of refed RFP cells with non-refed GFP cells (NF(GFP)/RF3(RFP), NF(GFP)/RF5(RFP)). The mixture of non-refed GFP cells and non-refed RFP cells (NF(GFP)/NF(RFP)) is the control. These results show that in the cell population of RF+NF, RF cells were more allocated to the solitary amoeboid cells than NF cells (GLMM and analysis of deviance, $P < 0.0001$, Table 3), and NF cells became more spore than RF cells (GLMM and analysis of deviance, $P < 0.0001$, Table 3). The GFP-RFP ratio was normalized by the ratio in NF(GFP)/NF(RFP). Sample size N represents three biological replicates with at least 3600 cells (amoeboid cells and spores) per condition.

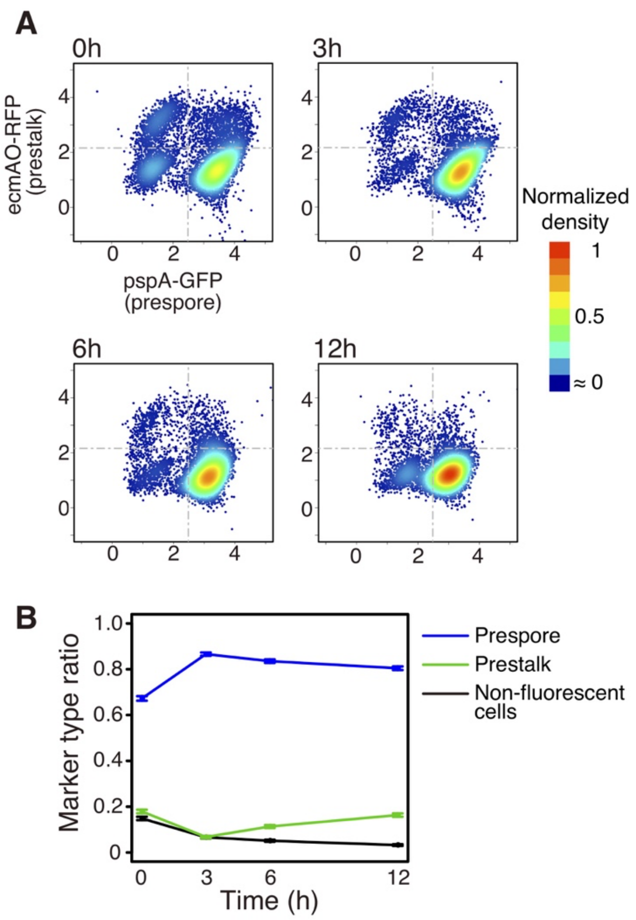


**Fig. S5. Flow cytometry analysis of cells under refeeding.**

(*A*) A scatter plot of fluorescent intensities of cell fate markers ecmAO-RFP (prestalk) and pspA-GFP (prespore) after 0, 3, 6, and 12h of incubation of dissociated slugs in the bacterial suspension. (*B*) Temporal change in the marker type ratio. Error bars indicate 95% confidence interval (CI). The ratio at the time of cell dissociation (0 h) was approximately prespore: prestalk: non-fluorescent cell = 0.6:0.2:0.2. Thus, in the model, the cell fate ratios of the full social commitment ($x = 1$) are designated as $SP_{initial} = 0.6$, $ST_{initial} = 0.2$, and $Sol_{initial} = 0.2$, respectively.

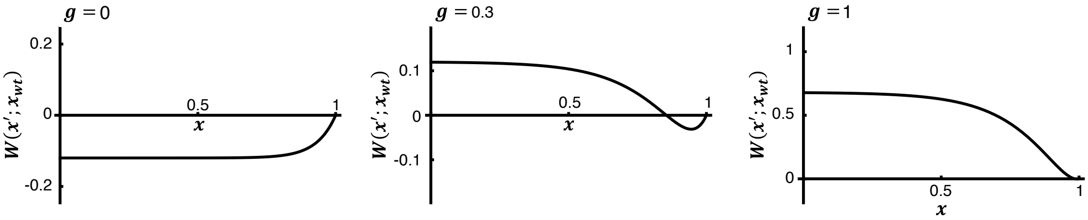


**Fig. S6. Alternative formulation of the invasion fitness as altruism.**

The invasion fitness of the solitary reversion mutant $W(x'; x_{wt})$ using the alternative formulation (Eq. **S15, S16**), which assumes explicit altruism: Success of sporulation depends on the average stalk investment in a group. The wild-type trait is $x_{wt} = 1$. $g$ is nutrient availability. These results were fundamentally the same result as invasion fitness with the equation Eq. **3** in the main text.

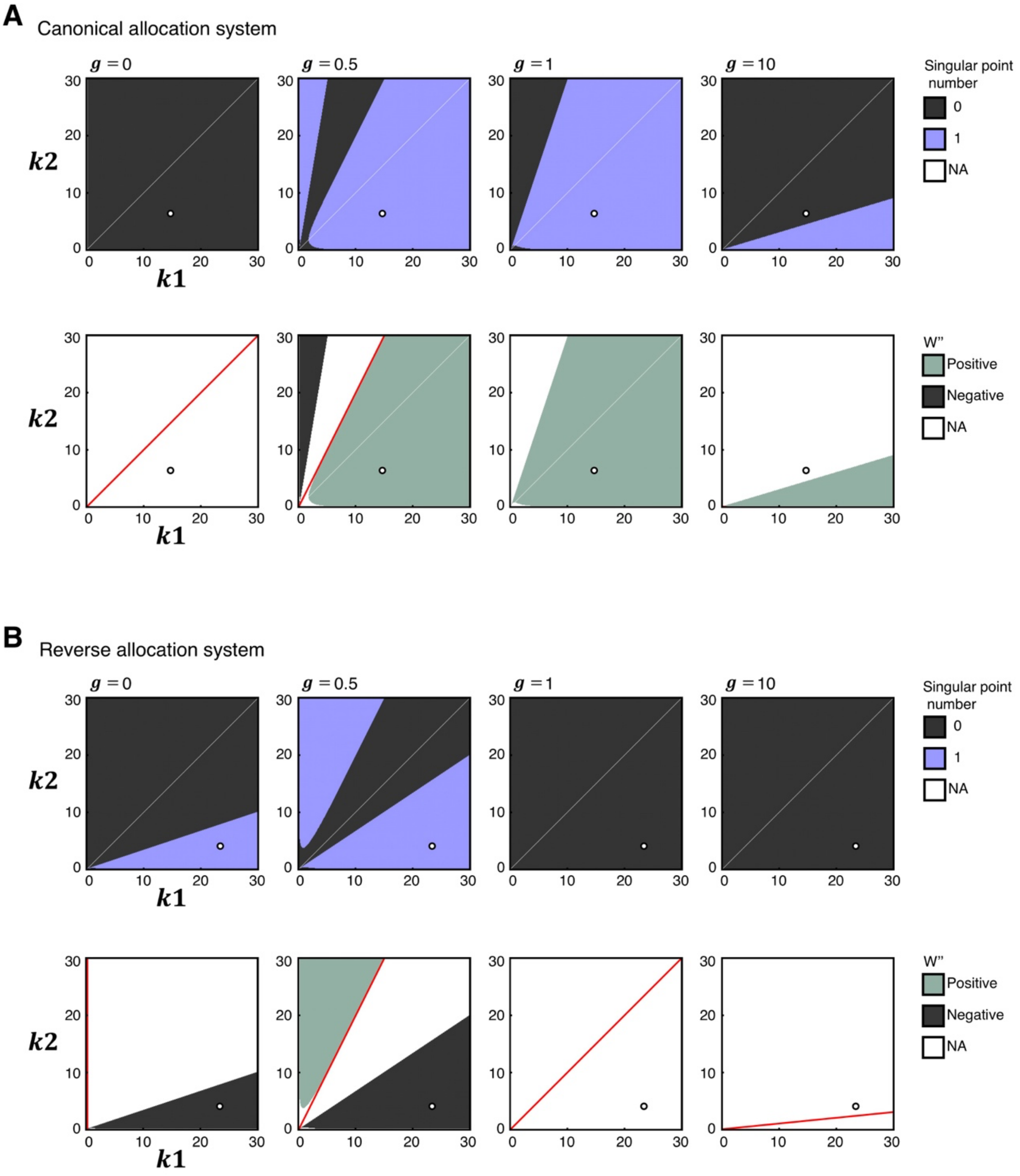


**Fig. S7. Parameter sensitivity to singularity and evolutionary conditions.**

(*A*) Change in $k1$, $k2$ and numerical calculation of the number of the singular point (Eq. **S4)** (upper panel) and the numerical calculation of the left-hand side of Eq. **S6** (second order differential of the fitness at the singular point, $W''$) (lower panel) in the canonical allocation system. The red lines indicate the analytical prediction describing the condition for ESS (Eq. **S10**). When $g < 1$, if $k1$, $k2$ are above the red line, the singular point is satisfied the condition for ESS. However, the condition for ESS is never satisfied, because we assume $k_1 \gg k_2$. When $g > 1$, in any $k1$, $k2$ ($k1 > 0$ and $k2 > 0$), the condition for ESS is also never satisfied. In addition, the criteria for convergence stable (Eq. **S7**) is also only satisfied under the same conditions as those described in the lower panel for ESS (see also Eq. **S8,** Eq. **S9**). Thus, the singular point is neither ESS nor convergence stable when evaluated under our assumed condition ($k_1 \gg k_2$).

(*B*) The number of the singular point and $W''$ in the reverse allocation system. When $g = 0$, if $0 < k1$, the singular point satisfied the condition for ESS. When $g < 1$, if $k1$, $k2$ is below the red line, the singular point satisfied the condition for ESS. In addition, the criteria for convergence stable (Eq. **S7**) is also satisfied under the same condition as shown in the lower panel for ESS (see also Eq. **S11,** Eq. **S12**). When $g = 1, 10$, there is no singular point in $0 \leq x \leq 1$. Thus, the singular point is evolutionally stable (ESS) and convergence stable under our assumed condition ($k_1 \gg k_2$). NA indicates that the value could not be calculated. White circles indicate the default value.

**Table S1.** Summary of statistical analysis. GLM is generalized linear model. $\beta_0$ , $\beta_1$ and $\beta_2$ indicate regression coefficients. Rv: Response variable, Ev: Explanatory variable. Link: Link functions that decide relationship between Rv and Ev. * indicates *P* <0.05

| **Analysis** | **Cell localization within aggregation for Figure 1*C*** |
|---|---|
| **Model formula**<br>**R code format**<br>**Explanation** | $g(d) = \beta_0 + \beta_1\, condition + \beta_2\, cellfate.$<br>$glm(d \sim condition + cellfate, family = gaussian(link = "identity")).$<br>GLM with gaussian error. Rv: $d$, Ev: $condition$ and $cellfate$. Link: $g(d) = d$.<br>$d$: The mean distance from center of a mound to each pixel. $condition$: Experimental condition, $cellfate$: Prespore or prestalk. |
| **Results** | **Overall model:** GLM and analysis of deviance, Cell fate: $F_{1,186}$ = 228.661, *P* < 0.0001*, Conditions: $F_{2,184}$ = 56.447, *P* < 0.0001*<br>**Multiple comparison:** $t$-test<br>1: Prespore (NF+NF) vs Prestalk (NF+NF): d.f. = 65.658, t = -14.222, adjusted *P* < 0.0001*<br>2: Prespore (NF+NF) vs Prespore (RF3+NF): d.f. = 65.472, t = -8.1747, adjusted *P* < 0.0001*<br>3: Prespore (NF+NF) vs Prestalk (RF3+NF): d.f. = 57.518, t = -16.525, adjusted *P* < 0.0001*<br>4: Prespore (NF+NF) vs Prespore (RF5+NF): d.f. = 36.687, t = -7.606, adjusted *P* < 0.0001*<br>5: Prespore (NF+NF) vs Prestalk (RF5+NF): d.f. = 39.133, t = -14.512, adjusted *P* < 0.0001*<br>6: Prestalk (NF+NF) vs Prespore (RF3+NF): d.f. = 63.852, t = 4.4839, adjusted *P* < 0.001*<br>7: Prestalk (NF+NF) vs Prestalk (RF3+NF): d.f. = 55.146, t = -6.1947, adjusted *P* < 0.0001*<br>8: Prestalk (NF+NF) vs Prespore (RF5+NF): d.f. = 35.066, t = 1.348, adjusted *P* = 0.373<br>9: Prestalk (NF+NF) vs Prestalk (RF5+NF): d.f. = 37.259, t = -4.973, adjusted *P* < 0.0001*<br>10: Prespore (RF3+NF) vs Prestalk (RF3+NF): d.f. = 63.301, t = -9.1856, adjusted *P* < 0.0001*<br>11: Prespore (RF3+NF) vs Prespore (RF5+NF): d.f. = 41.336, t = -1.7063, adjusted *P* = 0.286<br>12: Prespore (RF3+NF) vs Prestalk (RF5+NF): d.f. = 44.369, t = -7.856, adjusted *P* < 0.0001*<br>13: Prestalk (RF3+NF) vs Prespore (RF5+NF): d.f. = 50.541, t = 5.7015, adjusted *P* < 0.0001*<br>14: Prestalk (RF3+NF) vs Prestalk (RF5+NF): d.f. = 53.52, t = 0.60077, adjusted *P* < 0.551<br>15: Prespore (RF5+NF) vs Prestalk (RF5+NF): d.f. = 47.565, t = -4.8928, adjusted *P* < 0.0001* |
| **Analysis** | **Quantitative real-time polymerase chain reaction analysis for Figure 1*D***<br>Analysis to detect nutrient-specific changes |
| **Model formula**<br>**R code format**<br>**Explanation** | $g(q) = \beta_0 + \beta_1\, condition.$<br>$glm(q \sim condition, family = gaussian(link = "identity")).$<br>GLM with gaussian error. Rv: $q$, Ev: $condition$, Link: $g(q) = q$.<br>$q$: The levels of relative gene expression.<br>$condition$: Experimental condition. |
| **Results** | To understand cell-state change incurred by refeeding, following comparisons are considered.<br>**Multiple comparison:** GLM and analysis of deviance<br>No-nutrient versus Bacteria:<br>emcA: $F_{1,16}$ = 0.1013, adjusted P = 1<br>pspA: $F_{1,16}$ = 0.2646, adjusted P = 1<br>acaA: $F_{1,16}$ = 7.2018, adjusted P = 0.01631*<br>carA: $F_{1,16}$ = 15.289, adjusted P = 0.001247*<br>pdsA: $F_{1,16}$ = 14.177, adjusted P = 0.003385*<br>dscA: $F_{1,16}$ = 13.907, adjusted P = 0.001981*<br>tgrC: $F_{1,16}$ = 1.0826, adjusted P = 0.627171<br>cadA: $F_{1,16}$ = 0.1726, adjusted P = 1<br>csaA: $F_{1,16}$ = 0.1726, adjusted P = 1<br>No-nutrient versus Growth medium:<br>emcA: $F_{1,16}$ = 0.4288, adjusted P = 1<br>pspA: $F_{1,16}$ = 0.0279, adjusted P = 1<br>acaA: $F_{1,16}$ = 12.666, adjusted P = 0.005232*<br>carA: $F_{1,16}$ = 24.939, adjusted P = 0.000265*<br>pdsA: $F_{1,16}$ = 11.535, adjusted P = 0.003689*<br>dscA: $F_{1,16}$ = 16.156, adjusted P = 0.001981*<br>tgrC: $F_{1,16}$ = 0.5538, adjusted P = 0.627171<br>cadA: $F_{1,16}$ = 0.0767, adjusted P = 1<br>csaA: $F_{1,16}$ = 0.0767, adjusted P = 1 |

The continuation of Table S1.

| **Analysis** | **Quantitative real-time polymerase chain reaction analysis for Figure 1*D***<br>Analysis to detect temporal differences in the level of gene expression |
|---|---|
| **Explanation** | Pair-wise comparisons of levels of relative gene expression were performed between the sampling time points 0h and 5h with the bootstrap method. For the detail, see SI text, 'Quantitative real-time polymerase chain reaction analysis'. |
| **Results** | Bootstrap method<br>No-nutrient buffer 0h vs 5h<br>emcA: t = 2.243471, *P* = 0.02881<br>pspA: t = 2.570983, *P* = 0.02438**<br>acaA: t = -0.8896212, *P* = 0.21216<br>carA: t = -13.71146, *P* = 0.01272**<br>pdsA: t = -13.44524, *P* = 0.01206**<br>dscA: t = -0.9597171, *P* = 0.1883<br>tgrC: t = 4.070634, *P* = 0.01408**<br>cadA: t = 3.525237, *P* = 0.01524**<br>csaA: t = -4.731203 , *P* = 0.0121**<br>Bacteria 0h vs 5h<br>emcA: t = 6.490245, *P* = 0.01275**<br>pspA: t = 6.184117, *P* = 0.01264**<br>acaA: t = 1.342081, *P* = 0.12785<br>carA: t = 0.9833843, *P* = 0.18435<br>pdsA: t = 1.421683, *P* = 0.1052<br>dscA: t = 3.987957, *P* = 0.01547**<br>tgrC: t = 0.9550938, *P* = 0.19265<br>cadA: t = 1.30704, *P* = 0.13153<br>csaA: t = -2.370863, *P* = 0.03002<br>Growth medium 0h vs 5h<br>emcA: t = 5.054192, *P* = 0.01153**<br>pspA: t = 4.0807, *P* = 0.01505**<br>acaA: t = 6.217846, *P* = 0.01101**<br>carA: t = 5.886053, *P* = 0.0126**<br>pdsA: t = 1.348934, *P* = 0.11539<br>dscA: t = 2.10459, *P* = 0.03932<br>tgrC: t = 8.379845, *P* = 0.00888**<br>cadA: t = 2.570532, *P* = 0.02075**<br>csaA: t = 7.610438, *P* = 0.01291**<br>No-dissociation 0h vs 5h<br>emcA: t = -2.934227, *P* = 0.01913**<br>pspA: t = 1.194258, *P* = 0.14051<br>acaA: t = 0.8418263, *P* = 0.24907<br>carA: t = 1.136168, *P* = 0.15566<br>pdsA: t = 0.007316311, *P* = 0.4876<br>dscA: t = 4.799018, *P* = 0.01373**<br>tgrC: t = 0.08839139, *P* = 0.5383<br>cadA: t =1.56493, *P* = 0.07794<br>csaA: t = 0.1677567, *P* = 0.43407<br><br>** indicates *P* <0.025.<br>The *p*-value was the percentile rank of the observed *t*-values within bootstrap sampling data. For a two-sided test, *P* <0.025 was used for the significance. |

The continuation of Table S1.

| Analysis | **Number of single cells in aggregation assay for Figure 1*E* (left)** |
|---|---|
| **Model formula**<br>**R code format**<br>**Explanation** | $g(single) = \beta_0 + \beta_1\, condition.$<br>$glm(single \sim condition, family = poisson(link = "log")).$<br>GLM with poisson error. Rv: $single$, Ev: $condition$, Link: $g(single) = log(single)$. $single$: The number of single-state cells. $condition$: Experimental condition. |
| **Results** | To understand cell cohesion change incurred by refeeding, following comparisons are considered.<br>**Multiple comparison:** GLM and analysis of deviance<br>Buffer 5 h versus Bacteria 5 h<br>deviance = 2612.3, residual deviance = 424.93, d. f. = (1, 4), adjusted $P < 0.0001$*<br>Buffer 5 h versus Growth medium 5 h<br>deviance = 5920.9, residual deviance = 748.4, d. f. = (1, 4), adjusted $P < 0.0001$* |
| **Analysis** | **Proportion of single prespore cells in aggregation assay for Figure 1*E* (right)** |
| **Model formula**<br>**R code format**<br>**Explanation** | $g(P_{prespore}) = \beta_0 + \beta_1\, condition.$<br>$glm(cbind(prespore, prestalk) \sim condition, family = binomial(link = \text{"cloglog"})).$<br>GLM with binomial error. Rv: $P_{prespore}$, Ev: $condition$, Link: $g(P_{prespore}) = log(1 - log(1 - P_{prespore}))$. $P_{prespore}$: Probability that single state cells were expressed prespore marker. $prespore$: Prespore cell number. $prestalk$: Prestalk cell number. $condition$: Experimental condition. |
| **Results** | To understand effect of refeeding on prespore ratio in the unattached cells, following comparisons are considered.<br>**Multiple comparison:** GLM and analysis of deviance<br>Buffer+EDTA versus Buffer 5 h<br>deviance = 133.4, residual deviance = 9.438, d.f. = (1,4), adjusted $P < 0.0001$*<br>Buffer+EDTA versus Bacteria 5 h<br>deviance = 0.31927, residual deviance = 13.964, d.f. = (1, 4), adjusted $P = 0.572$<br>Buffer+EDTA versus Growth medium 5 h<br>deviance = 7.228, residual deviance = 17.483, d.f. = (1,4), adjusted $P = 0.01435508$*<br>Buffer 5 h versus Bacteria 5 h<br>deviance = 131.66, residual deviance = 9.073, d.f. = (1,4), adjusted $P < 0.0001$*<br>Buffer 5 h versus Growth medium 5 h<br>deviance = 159.01, residual deviance = 12.592, d.f. = (1,4), adjusted $P < 0.0001$* |
| **Analysis** | **Cell localization in an upper part of a fruiting body for Figure 2*B*.** |
| **Model formula**<br>**R code format**<br>**Explanation** | $g(fspore) = \beta_0 + \beta_1\, fstalk.$<br>$glm(fspore \sim fstalk, family = Gamma(link = "log")).$<br>GLM with gamma error. Rv: $fspore$, Ev: $fstalk$. Link: $g(fspore) = log(fspore)$.<br>$fspore$: Prespore frequency at each vertical position in a fruiting body. $fstalk$: Prestalk frequency at the corresponding vertical position. To use log link function, we added 0.1 to the score of $fspore$ and $fstalk$ before analysis, because these included a score 0. |
| **Results** | No-dissociation: slope =-4.69333 ± 0.28704, $P < 0.0001$*<br>NF+NF: slope = -2.03408 ± 0.26450, $P < 0.0001$*<br>RF3+NF: slope = 4.52924 ± 0.24770, $P < 0.0001$*<br>RF5+NF: slope =3.69829 ± 0.30505, $P < 0.0001$* |
| **Analysis** | **Prespore marker expression ratio in a basal disk for Figure 2*E*.** |
| **Model formula**<br>**R code format**<br><br>**Explanation** | $g(Iprespore) = \beta_0 + \beta_1\, condition + log(Iprestalk).$<br>$glm(Iprespore \sim condition, offset = log(Iprestalk), family = gaussian(link = "identity")).$<br>GLM with gaussian error. Rv: $Iprespore$, Ev: $condition$, Link: $g(Iprespore) = Iprespore$. $Iprespore$: Intensity value of prespore cell marker expression, $Iprestalk$: Intensity value of prestalk marker expression, $condition$: Experimental condition. $log(Iprestalk)$ is an offset term to adjust $Iprespore$ toward $Iprestalk$. |
| **Results** | **Overall model:** GLM and analysis of deviance, Conditions: $F_{3,94} = 25.008$, $P < 0.0001$*<br>**Multiple comparison:** $t$-test<br>NF+NF versus No-dissociation: d.f. = 53.998, t = 1.9158, adjusted $P = 0.06069$<br>NF+NF versus RF3+NF: d.f. = 22.468, t = -9.1623, adjusted $P < 0.0001$*<br>NF+NF versus RF5+NF: d.f. = 18.888, t = -12.292, adjusted $P < 0.0001$* |

The continuation of Table S1.

| | |
|---|---|
| **Analysis** | **Prespore ratio among cells left behind in basal region for Figure 2G.** |
| **Explanation** | Same as the model formula for "Prespore marker expression ratio in a basal disk for Fig. 2E". |
| **Results** | **Overall model:** GLM and analysis of deviance, Conditions: F3,86 = 41.432, P < 0.0001*<br>**Multiple comparison:** *t*-test<br>NF+NF versus No-dissociation: d.f. = 43.841, t = 3.7784, adjusted *P* < 0.001*<br>NF+NF versus RF3+NF: d.f. = 23.08, t = -7.2017, adjusted *P* < 0.0001*<br>NF+NF versus RF5+NF: d.f. = 23.099, t = -21.81, adjusted P < 0.0001* |
| **Analysis** | **Growth cost of the social commitment for Figure 3*B*.**<br>Comparison between cell growth with and without the social commitment |
| **Model formula**<br>**R code format**<br>**Explanation** | $g(cellnumber) = \beta_0 + \beta_1\, condition + \beta_2\, day.$<br>$glm(cellnumber \sim condition + day, family = gaussian(link = "identity")).$<br>GLM with gaussian error. Rv: $cellnumber$, Ev: $condition$ and $day$.<br>Link: $g\ (cellnumber\ ) = cellnumber$ . $cellnumber$ : Logarithmic transformed cell number. $condition$: Experimental condition, $day$ : Day after starting the growth assay. |
| **Results** | To understand effect of social commitment on cell growth, following comparisons are considered.<br>GLM and analysis of deviance,<br>'Refed & No-dissociation' versus 'Refed & Dissociation': $F_{1,21}$ = 12.175, *P* = 0.002187* |
| **Analysis** | **Growth cost of the social commitment for Figure 3*B*.**<br>Growth curve fitting. |
| **Model formula**<br><br>**R code format**<br>**Explanation** | $y = \frac{D}{(1+\exp\,(\frac{4j}{D}(lag-t)+2))}.$<br>$\mathrm{nls(y \sim D/(1 + exp}\left(\left(4 * \frac{\mathrm{j}}{\mathrm{D}}\right) * (lag - \mathrm{t}) + 2\right))).$<br>$y$: the cell number, $D$: maximum cell number, $t$: time. $j$: growth rate, $lag$: length of the lag phase. |
| **Results** | Nonlinear least squares method<br>Estimated growth rate $j$<br>'Refed & Dissociation': 0.067 ± 0.012, 'Refed & No-dissociation': 0.064 ± 0.012, 'Continuous starvation': 0.057 ± 0.009.<br>Estimated maximum cell number $D$<br>'Refed & Dissociation': 2.69758 ± 0.10125, 'Refed & No-dissociation': 2.80108 ± 0.09965, 'Continuous starvation': 2.688191 ± 0.117792.<br>Estimated lag time $lag$<br>'Refed & Dissociation': 15.24 ± 4.27 h, 'Refed & No-dissociation': 23.97 ± 4.35 h, 'Continuous starvation': 36.17 ± 4.03 h. |
| **Analysis** | **Allocation between social and solitary fitness for Figure S4.** |
| **Model formula**<br>**R code format**<br><br>**Explanation** | $g(P_{gfp}) = \beta_0 + \beta_1\, condition + log(P_{initial}) + r.$<br>$glmer(cbind(gfp, rfp) \sim condition + (1\|r), offset = log(P_{initial}), family = binomial(link = "cloglog")).$<br>GLMM with binomial error. Rv: $P_{gfp}$, Ev: $condition$, Link: $g(P_{gfp}) = log(1 - log(1 - P_{gfp}))$. $P_{gfp}$: Probability of cells (or spores) expressed constitutive GFP in terminal differentiation. $gfp$: GFP cell number, $rfp$: RFP cell number, $condition$: Experimental condition, $P_{initial}$: Ratio of cells with constitutive GFP toward RFP in cells of immediately after mixing. $log(P_{initial})$ is an offset term to adjust $P_{gfp}$ by $P_{initial}$. $r$ is the difference by replicate experiments as a random effect. |
| **Results** | GLMM and analysis of deviance<br>For Figure S4, *left,*<br>Conditions: (NF(GFP)/NF(RFP)), (RF3(GFP)/NF(RFP)), (RF5(GFP)/NF(RFP))<br>Effect of condition on probability of GFP cells, deviance = 189.41, d.f. = (4, 2), *P* < 0.0001*<br>Effect of condition on probability of GFP spores, deviance = 368.41, d.f. = (4, 2), *P* < 0.0001*<br><br>For Figure S4, *Right,*<br>Conditions: (NF(GFP)/NF(RFP)), (NF(GFP)/RF3(RFP)), (NF(GFP)/RF5(RFP))<br>Effect of condition on probability of GFP cells, deviance = 133.32, d.f. = (4,2), *P* <0.0001*<br>Effect of condition on probability of GFP spores, deviance = 336.84, d.f. = (4,2), *P* <0.0001* |

**Table S2.** Primer pairs and fluorescent beacons for qRT-PCR. * indicates primers and beacons that are newly constructed in this study. The rest are previously described [19].

| Gene name | | 5' →3' | Location | | Reporter Dye |
|---|---|---|---|---|---|
| *rnlA* | Forward | CGGATAAAAGGTACGCTAGGGATA | 2327 | 2350 | - |
| | Reverse | GTGCCGAACCACATAACAGATATG | 2375 | 2398 | - |
| | Probe | CAGGCTAGTCACATATT | 2352 | 2368 | VIC |
| *acaA* | Forward | TTGGTATTAGTCATGGTCCTTTGG | 3833 | 3856 | - |
| | Reverse | GAGGCGGTATTGGCAGTATCA | 3903 | 3923 | - |
| | Probe | CTGGTTGTATCGGTATCAG | 3860 | 3878 | FAM |
| *carA* | Forward | AAATATGTTTCCACCAGCACTCAA | 693 | 716 | - |
| | Reverse | GATAAATGTGACAGATGCCCAAAA | 751 | 774 | - |
| | Probe | ATTCTCCACACCTATTTG | 718 | 735 | FAM |
| *pdsA* | Forward | AGCAAGTGGCATTGAATATCCA | 789 | 807 | - |
| | Reverse | ACCAAAGACATAGTGGTGGCATT | 829 | 851 | - |
| | Probe | TCACAGAGTTGGTCCC | 809 | 824 | FAM |
| *tgrC1* | Forward | CCTCCAACACCAATAGATGCAA | 64 | 85 | - |
| | Reverse | GTTCTGGGTTCTTTTTCGTTTTTATACA | 152 | 179 | - |
| | Probe | TAATAGTAATCTCCCATATTCTACC | 117 | 141 | FAM |
| *ecmA* | Forward | GTTAATGCGGAAACTGAAACCA | 58 | 79 | - |
| | Reverse | CAAAAAGTAAACCTGCAGAACACA | 146 | 169 | - |
| | Probe | ACAAACCAATACAGCATGTG | 81 | 100 | FAM |
| *pspA* | Forward | GCGCTGATCAAACTTCTTCACAT | 263 | 285 | - |
| | Reverse | GGGTGTGGCAGTGATTTTACAA | 312 | 333 | - |
| | Probe | CACTCGGTTCTGATTGG | 287 | 303 | FAM |
| *dscA* | Forward | GGTCGTGGTGATGCTGATCA | 232 | 251 | - |
| | Reverse | CGATATTCAAACCAGGAAACATTATC | 286 | 311 | - |
| | Probe | TACATCATACAAAATCCG | 258 | 275 | FAM |
| *cadA* * | Forward | AATTGGCTCAAGGCAGTACAAAC | 209 | 231 | - |
| | Reverse | AAAAGCTCCTGGTAAGACTTGGAA | 265 | 288 | - |
| | Probe | TAACCTCAATAAATGGTCTTTC | 239 | 260 | FAM |
| *csaA* * | Forward | AACAATTTATTTCCTCGTGCCAAA | 462 | 485 | - |
| | Reverse | TTGAAAAGCCAAATGGTTGAATG | 519 | 541 | - |
| | Probe | CAATCGCTGGTGGTCTA | 488 | 504 | FAM |

**Table S3.** Representative variables and parameters for the model in Figure 4,5.

| Notation | Description |
|---|---|
| $x$ | A degree of association with other cells. An evolutionary trait. |
| $x_{wt}$ | The trait $x$ of the wild-type cell. The default value is $x_{wt} = 1$ indicating full social commission. |
| $x'$ | Trait $x$ of the rare mutant cell. |
| $P(SP\|x)$ | The probability of a cell with the trait $x$ to be a prespore cell. |
| $P(ST\|x)$ | The probability of a cell with the trait $x$ to be a prestalk cell. |
| $P(Sol\|x)$ | The probability of a cell with the trait $x$ to be a solitary cell. |
| $SP_{initial}$ | The probability to be a prespore cell at the initial state of the evolution ($x = 1$). The default value is 0.6. |
| $ST_{initial}$ | The probability to be a prestalk cell at the initial state of the evolution. The default value is 0.2. |
| $Sol_{initial}$ | The probability to be a solitary cell at the initial state of the evolution. The default value is 0.2. |
| $f(x'; x_{wt})$ | The survival probability of the mutant cell with $x'$ when the mutant cell interacts with the wild-type cell. Because the mutant is rare, the interaction between mutant cells is ignored. |
| $f(x_{wt}; x_{wt})$ | The survival probability of the wild-type cell with $x_{wt}$ when the wild-type cell interacts with another wild-type cell. |
| $W(x'; x_{wt})$ | Invasion fitness of the mutant with $x'$ when the mutant cell interacts with the wild-type cell. |
| $g$ | The additional weight to the solitary fitness as defined by nutrient availability. |
| $k_1, k_2$ | The rates of cell-fate transition. |
| $i$ | Additional spore investment. An evolutionary trait. |
| $i_{wt}$ | The trait $i$ of the wild-type cell. The default value is 0. |
| $i'$ | The trait $i$ of the cheating mutant cell. |
| $P_t(SP\|i)$ | The probability of a cell with $i$ to be a prespore cell at the time $t$ |
| $P_t(ST\|i)$ | The probability of a cell with $i$ to be a prestalk cell at the time $t$ |
| $P_t(Sol\|i)$ | The probability of a cell with $i$ to be a solitary cell at the time $t$ |
| $f(i'; i_{wt})$ | The survival probability of the mutant cell with $i'$when the mutant cell interacts with the wild-type cell. |
| $f(i_{wt}; i_{wt})$ | The survival probability of the wild-type cell with $i_{wt}$ when the wild-type cell interacts with another wild-type cell. |
| $W(i'; i_{wt})$ | Invasion fitness of the cheating mutant with $i'$when the mutant cell interacts with the wild-type cell. |

**Movie S1 (separate file).** The process of aggregation formation in NF+NF condition. Images were obtained every 12 s. For the explanation, see Figure S1.

**Movie S2 (separate file).** The process of aggregation formation in RF3+NF condition. Images were obtained every 12 s. For the explanation, see Figure S1.

**Movie S3 (separate file).** The process of fruiting body formation in NF+NF condition. Images were obtained every 3 min. For the explanation, see Figure S3*B*.

**Movie S4 (separate file).** The process of fruiting body formation in RF3+NF condition. Images were obtained every 3 min. For the explanation, see Figure S3*B*.

**Movie S5 (separate file).** Solitary cells left behind in the base of the fruiting body in RF5+NF condition. Some moving cells were observed in the base 7 h after plating. Images were obtained every 10 s. The numbers on the upper right side indicate min:sec. Scale bars = 50 µm.

**Dataset S1 (separate file).** Data for the experiments.

**Code S1 (separate file).** A Mathematica 10.3 code for the theoretical models.

#### SI References


[1] Sawai S, Guan XJ, Kuspa A, Cox EC. High-throughput analysis of spatio-temporal dynamics in *Dictyostelium*. Genome Biology. 2007;8(7):R144. doi:10.1186/gb-2007-8-7-r144

[2] Taniguchi D, Ishihara S, Oonuki T, Honda-Kitahara M, Kaneko K, Sawai S. Phase geometries of two-dimensional excitable waves govern self-organized morphodynamics of amoeboid cells. Proceedings of the National Academy of Sciences of the United States of America. 2013;110(13):5016–5021. doi:10.1073/pnas.1218025110

[3] Dingermann T, Reindl N, Werner H, Hildebrandt M, Nellen W, Harwood A, Williams J, Nerke K. Optimization and in situ detection of Escherichia coli β-galactosidase gene expression in *Dictyostelium discoideum*. Gene. 1989;85(2):353–362. doi:10.1016/0378-1119(89)90428-9

[4] Fey P, Compton K, Cox EC. Green fluorescent protein production in the cellular slime molds *Polysphondylium pallidum* and *Dictyostelium discoideum*. Gene. 1995;165(1):127–130.

[5] Masaki N, Fujimoto K, Honda-Kitahara M, Hada E, Sawai S. Robustness of self-organizing chemoattractant field arising from precise pulse induction of its breakdown enzyme: A single-cell level analysis of PDE expression in *Dictyostelium*. Biophysical Journal. 2013;104(5):1191–1202. doi:10.1016/j.bpj.2013.01.023

[6] Abràmoff MD, Magalhães PJ, Ram SJ. Image Processing with ImageJ. Biophotonics International. 2004;11:36–42.

[7] R: A language and environment for statistical computing. R Foundation for Statistical Computing, Vienna, Austria. URL https://www.R-project.org/. 2015.

[8] Harrington BJ, Raper KB. Use of a fluorescent brightener to demonstrate cellulose in the cellular slime molds. Applied Microbiology. 1968;16(1):106–113.

[9] Efron B, Tibshirani R. An introduction to the bootstrap. Chapman & Hall; 1993. (Chapman & Hall).

[10] Parkinson K, Bolourani P, Traynor D, Aldren NL, Kay RR, Weeks G, Thompson CRL. Regulation of Rap1 activity is required for differential adhesion, cell-type patterning and morphogenesis in *Dictyostelium*. Journal of Cell Science. 2009;122(3):335–344. doi:10.1242/jcs.036822

[11] Zwietering MH, Jongenburger I, Rombouts FM, van't Riet K. Modeling of the bacterial growth curve. Applied and Environmental Microbiology. 1990;56(6):1875–1881. doi:10.1128/aem.56.6.1875-1881.1990

[12] Geritz SAH, Kisdi É, Meszéna G, Metz JAJ. Evolutionarily singular strategies and the adaptive growth and branching of the evolutionary tree. Evolutionary Ecology. 1998;12(1):35–57. doi:10.1023/a:1006554906681

[13] Dieckmann U, Ferrière R. Adaptive Dynamics and Evolving Biodiversity. In: Ferrière R, Dieckmann U, Couvet D, editors. Evolutionary conservation biology. Cambridge, UK: Cambridge University Press; 2004. p. 188–224.

[14] Smith JM. Evolution and the theory of games. Cambridge University Press; 1982. (Cambridge University Press).

[15] Eshel I. Evolutionary and continuous stability. Journal of Theoretical Biology. 1983;103(1):99–111. doi:10.1016/0022-5193(83)90201-1

[16] Dubravcic D, Baalen M van, Nizak C. An evolutionarily significant unicellular strategy in response to starvation in *Dictyostelium* social amoebae. F1000Research. 2014;3:133. doi:10.12688/f1000research.4218.2

[17] Sternfeld J. A study of PstB cells during *Dictyostelium* migration and culmination reveals a unidirectional cell type conversion process. Roux's archives of developmental biology. 1992;201(6):354–363. doi:10.1007/bf00365123

[18] Abe T, Early A, Siegert F, Weijer C, Williams J. Patterns of cell movement within the *Dictyostelium* slug revealed by cell type-specific, surface labeling of living cells. Cell. 1994;77(5):687–699.

[19] McQuade KJ, Nakajima A, Ilacqua AN, Shimada N, Sawai S. The green tea catechin epigallocatechin gallate (EGCG) blocks cell motility, chemotaxis and development in *Dictyostelium discoideum* Harwood AJ, editor. PLoS ONE. 2013;8(3):e59275. doi:10.1371/journal.pone.0059275
